# Synthetic coevolution reveals adaptive mutational trajectories of neutralizing antibodies and SARS-CoV-2

**DOI:** 10.1101/2024.03.28.587189

**Authors:** Roy A. Ehling, Mason Minot, Max D. Overath, Daniel J. Sheward, Jiami Han, Beichen Gao, Joseph M. Taft, Margarita Pertseva, Cédric R. Weber, Lester Frei, Thomas Bikias, Ben Murrell, Sai T. Reddy

**Author notes:** Equal contribution (Roy A. Ehling, Mason Minot).

## Abstract

The Covid-19 pandemic showcases a coevolutionary race between the human immune system and SARS-CoV-2, mirroring the Red Queen hypothesis of evolutionary biology. The immune system generates neutralizing antibodies targeting the SARS-CoV-2 spike protein’s receptor binding domain (RBD), crucial for host cell invasion, while the virus evolves to evade antibody recognition. Here, we establish a synthetic coevolution system combining high-throughput screening of antibody and RBD variant libraries with protein mutagenesis, surface display, and deep sequencing. Additionally, we train a protein language machine learning model that predicts antibody escape to RBD variants. Synthetic coevolution reveals antagonistic and compensatory mutational trajectories of neutralizing antibodies and SARS-CoV-2 variants, enhancing the understanding of this evolutionary conflict.

## INTRODUCTION

The Covid-19 pandemic triggered a dynamic and ongoing interaction between the human immune system and SARS-CoV-2. On one hand, humans produce neutralizing antibodies that target the viral spike protein receptor binding domain (RBD), which is essential for viral entry into host cells. On the other hand, SARS-CoV-2 is able to mutate its RBD to evade antibody recognition and increase its infectivity. Importantly, widespread immunity within populations, resulting from vaccination or natural infection, exerts selective pressure on the virus, driving the emergence of escape mutations that can evade the existing neutralizing antibodies (*1*). This leads to a coevolutionary arms race between antibodies and SARS-CoV-2, where each side tries to gain an advantage over the other through mutational adaptation. For example, after encountering a new variant such as Omicron BA.1, pre-existing memory B cells that are specific to an earlier variant (e.g., ancestral Wu-Hu-1) can undergo somatic hypermutation to generate antibodies with enhanced binding and neutralization to Omicron BA.1 (*2*, *3*). Conversely, SARS-CoV-2 can evolve variants with RBD mutations that reduce or escape binding and neutralization to existing antibodies (*4–6*). This process of antagonistic coevolution resembles the Red Queen hypothesis of evolutionary biology (*7*). Here, we establish a synthetic coevolution system that employs high-throughput screening of antibody and RBD variant libraries using protein mutagenesis, cell surface display and deep sequencing; additionally we train and implement a protein language machine learning model capable of accurately predicting antibody escape to RBD variants (**Fig. 1**). We apply synthetic coevolution to uncover the antagonistic and compensatory mutational trajectories of neutralizing antibodies and SARS-CoV-2 variants.

**Figure 1.**
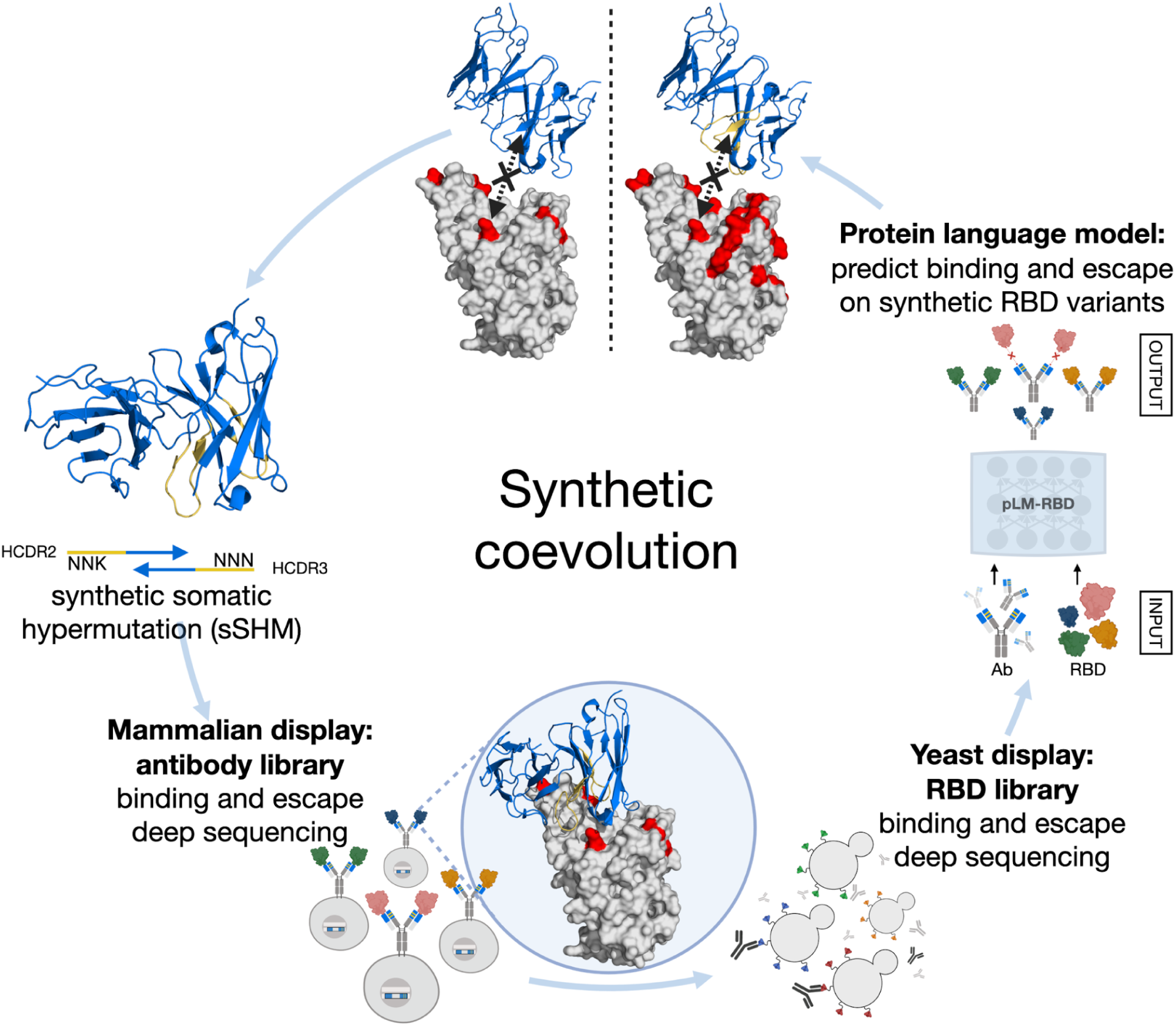
Overview of synthetic coevolution system for neutralizing antibodies and SARS-CoV-2 RBD. Antibodies that fail (blue ribbon) to bind an RBD variant (gray: RBD, red: mutations) undergo synthetic somatic hypermutation (sSHM) through targeted mutagenesis of the HDR2 and HDR3 (yellow). Antibody clonotype (sSHM) libraries are screened by mammalian surface display to select for clones that recover binding to RBD variants. The variant-selected antibodies and clonotype libraries are screened for binding and escape to synthetic RBD variants by yeast surface display. The RBD sequences for binding and escape fractions are determined by deep sequencing and used to train a protein language machine learning model, which can predict antibody binding and escape across a large mutational landscape of SARS-CoV-2.

## RESULTS

### Evolving neutralizing antibodies for binding to SARS-CoV-2 variants by synthetic somatic hypermutation

Eight neutralizing antibodies were selected representing the three major binding classes (Class 1-3) (*8*) that directly engage with the ACE2-binding site of the RBD (**Fig. 2A, Table S1**). Also, several of the antibodies possessed features associated with convergent germlines or public clones (*2*, *9*), suggesting they may be representative of antibodies that can shape viral evolution. The selected antibodies possessed high reactivity to the RBD of the ancestral Wu-Hu-1 and varying degrees of cross-reactivity to other variants, with most having substantially reduced reactivity to Omicron BA.1 (BA.1) (*4*). In order to mimic naturally occurring somatic hypermutation of antibodies, from each precursor (wild-type) antibody clone, we generated a library of clonal variants (clonotypes) through a process we refer to as synthetic somatic hypermutation (sSHM). Antibody clonotype libraries were designed to incorporate two degenerate NNK codons (N=A, G, C, T; K = G or T) in the variable heavy chain (V_H_) complementarity determining region 2 (HCDR2) and 3 (HCDR3) (**Fig. 2B**). Given a typical length of HCDR2 (4-6 a.a.) and HCDR3 (10-20 a.a.), the theoretical diversity of clonotype libraries ranged from 16,000 - 48,000 variants. Clonotype libraries were generated by PCR mutagenesis and cloned into homology directed repair (HDR) templates for CRISPR-Cas9 integration into an established mammalian cell antibody display and secretion platform (plug-and-(dis)play, PnP cells) (**Fig. S1**) (*10*, *11*).

**Figure 2.**
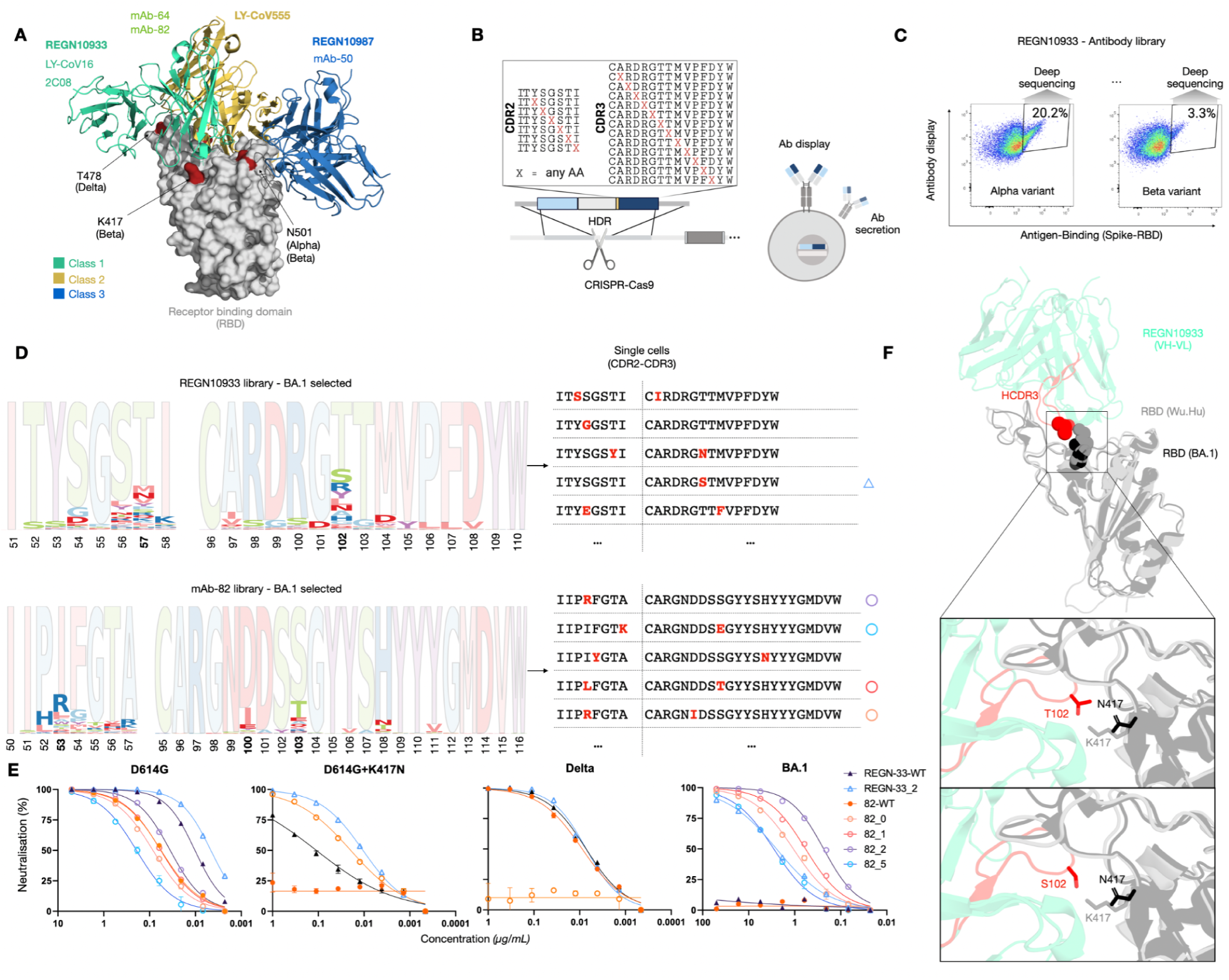
Recovery of antibody binding and neutralization by synthetic somatic hypermutation (sSHM). (**A**) Crystal structure of selected antibodies (bold) and corresponding epitope classifications shown in cyan (Class 1), yellow (Class 2) and blue (Class 3), with key mutations in the Alpha, Beta and Delta strains highlighted in red. (**B**) Representative mutagenesis scheme for sSHM, targeting the antibody variable heavy chain HCDR2 and HCDR3 with one mutation each (X), followed by the genomic integration of the library into a mammalian cell antibody display and secretion platform by CRISPR-Cas9 and homology-directed repair (HDR). (**C**) Representative FACS plots for REGN10933 antibody clonotype (sSHM) libraries screened for binding to Alpha and Beta RBD variants. Selected populations are sorted and deep sequenced. (**D**) Protein sequence logos of antibody clonotype pools (HCDR2 and HCDR3) following selection for binding to Omicron BA.1; opaque residues correspond to WT antibody sequence (left). The sequences of single antibody clones from the BA.1-binding population; mutated positions of HCDR2 and HCDR3 are depicted in red (right). (**E**) Viral neutralization with monoclonal antibodies from BA.1-selected clonotype libraries shown in (D), reduction in luciferase signal indicates successful viral neutralization. (**F**) Superimposed crystal structure of WuHu (PBD: 6XDG_A) and Omicron BA.1 (PDB: 7WRZ) with REGN10933 (PDB: 6XDG_C).

Antibody libraries in PnP cells were screened by fluorescence-activated cell sorting (FACS) to isolate clonotypes that possessed binding to RBD variants (Alpha, Beta, Delta or BA.1) (**Fig. 2C, Fig. S2**). The sSHM libraries LY-CoV16_sSHM_ and LY-CoV555_sSHM_ produced binders for only Alpha and Delta variants (**Fig. S2, S3**). The precursor LY-CoV16_WT_ has 100-fold reduced neutralization against Alpha (*12*), which may explain this lower binding compared to REGN10933_sSHM_, REGN10987_sSHM_ and LY-CoV555_sSHM_ libraries (**Fig. S2C**). LY-CoV555_WT_ loses nearly all binding and neutralization activity against the Delta variant (*12*), however LY-CoV555_sSHM_ possessed clonotypes with specificity to Delta (**Fig. S3**). The mAb-82_sSHM_ but not mAb-64_sSHM_ resulted in a complete loss of binding to the Delta variant (slightly reduced affinity of mAb-82_WT_), while both libraries could recover clonotypes with binding to the Beta variant, their wildtype versions have extremely low affinity to Beta, most likely due to the K417N mutation (*10*). Relative to their precursor antibodies, the class 1 REGN10933_sSHM_ and class 1-like mAb-82_sSHM_ and mAb-64_sSHM_ libraries possessed clonotypes with substantially improved binding to BA.1 (**Fig. S4**). This suggests that to regain specificity for BA.1, class 1 and class 1-like antibodies must overcome dependency on K417 or adapt to K417N, which is consistent with reported results (*13*). Neither mAb-50_sSHM_ nor REGN10987_sSHM_ possessed clonotypes that recovered significant binding to BA.1, this is likely because they were unable to overcome the class 3 epitope mutations in BA.1 (N440K and G446S) (*5*). The antibody 2C08_WT_ has been previously shown to have broad specificity and neutralization to most SARS-CoV-2 variants (*9*), therefore, expectedly, the corresponding 2C08_sSHM_ recovered clonotypes with binding to all variants including BA.1 (**Fig. S2, S4**).

Following selection of clonotypes with specificity to the different RBD variants, genomic DNA was extracted from PnP cells and targeted deep sequencing was performed on the V_H_ region. Sequencing analysis revealed both considerable diversity as well as conserved positions across most clonotypes (**Fig. S3**). The reliance of many antibodies on public or convergent germlines that are associated with binding to SARS-CoV-2 (*2*, *9*) may explain these limitations.

Across the variant-selected sSHM libraries, with few exceptions, there was more conservation in HCDR3 versus HCDR2 (**Fig. S3**). Within the HCDR3 of BA.1-selected REGN10933_sSHM_, mutations in position T102 appear to be sufficient to overcome BA.1 escape (**Fig. 2D**). For mAb-82_sSHM_, positions D100, S103 as well as position I53 exhibit the largest degree of diversity (**Fig. 2D**). The crystal structure of REGN10933 in complex with the RBD from Wu-Hu-1 or BA.1 suggests that the T102S mutation in HCDR3 interacts with the K417N mutation of the RBD (**Fig. 2E**). At 102 of HCDR3, the threonine is mutated to serine, which lacks the methyl substituent at the Beta-carbon but broadly shares the same physicochemical properties (similar pK, hydroxyl-group at Beta-carbon), thus a clash or interference of the methyl-group with N417 but not K417 is implied. The disruption of the salt bridge between D30 of ACE2 and K417 has been shown to reduce affinity and infectivity of SARS-CoV-2. Nevertheless, mutations in position 417 have been identified in multiple lineages (Beta, Gamma, BA.1) and thus appear to be one of the drivers behind escape of both Beta and BA.1 from REGN10933.

In addition to deep sequencing, PnP hybridoma cells from clonal libraries of REGN10933_sSHM_ and mAb-82_sSHM_ were single-cell sorted and expanded, followed by purification of antibody proteins from cell supernatant (*11*). With this subset of antibodies, we performed neutralization assays using HEK293T-ACE2 cells and SARS-CoV-2 pseudotyped lentivirus (*14*). While all tested wild-type antibodies and antibody clones maintained neutralization to the ancestral SARS-CoV-2 (D614G), only the sSHM-derived clones were able to neutralize the BA.1 virus (**Fig. 2F**). Interestingly, mAb-82 clones showed no significant neutralization against the Delta variant even at the highest concentration tested (1 ug/ml) compared to mAb-82_WT_ (IC_50_ = 0.0088 µg/mL), confirming the lack of Delta binding observed during library selection (**Fig. S2**). While it has been observed that affinity maturation generally increases antibody breadth and binding affinity to variants (*2*, *15*, *16*), it is conceivable that certain adaptations may produce local maxima that prevent further breadth.

### Coevolution of RBD variants and their mutational mapping to neutralizing antibodies

Next, we comprehensively mapped the specificity of sSHM-derived antibodies and their clonal libraries to identify potential RBD escape mutations and variants. First, we performed mutational mapping by integrating yeast display screening of RBD mutagenesis libraries for binding or escape (non-binding) to ACE2 or neutralizing antibodies, followed by deep sequencing. As described previously (*17*), we used tiling RBD mutagenesis libraries (library RBM-1T, RBM-2T and RBM-3T) targeting the core regions of the receptor-binding motifs (RBM-3: positions 439–452; RBM-1: 453–478; RBM-2: 484–505), which are subregions of the RBD that directly interface with ACE2 and correspond to epitope sites of class 1, 2 and 3 neutralizing antibodies. Additional libraries were constructed consisting of three degenerate NNK codons tiled across RBM-2 and possessing the dominant K417N and observed K417T mutation (library RBM-2_K417N/T_) or across RBM-1 containing the frequently observed mutations L452R (Delta, BA.4/5) and T478K (Delta, BA.1/2/3/4/5) (library RBM-1_L452R+T478K_) (**Fig. 3A**, **Fig. S5**). This resulted in library sizes of 4.59 x 10^6^ for RBM-2_K417N/T_ and and 0.96 x 10^6^ for RBM-1_L452R+T478K_.

**Figure 3.**
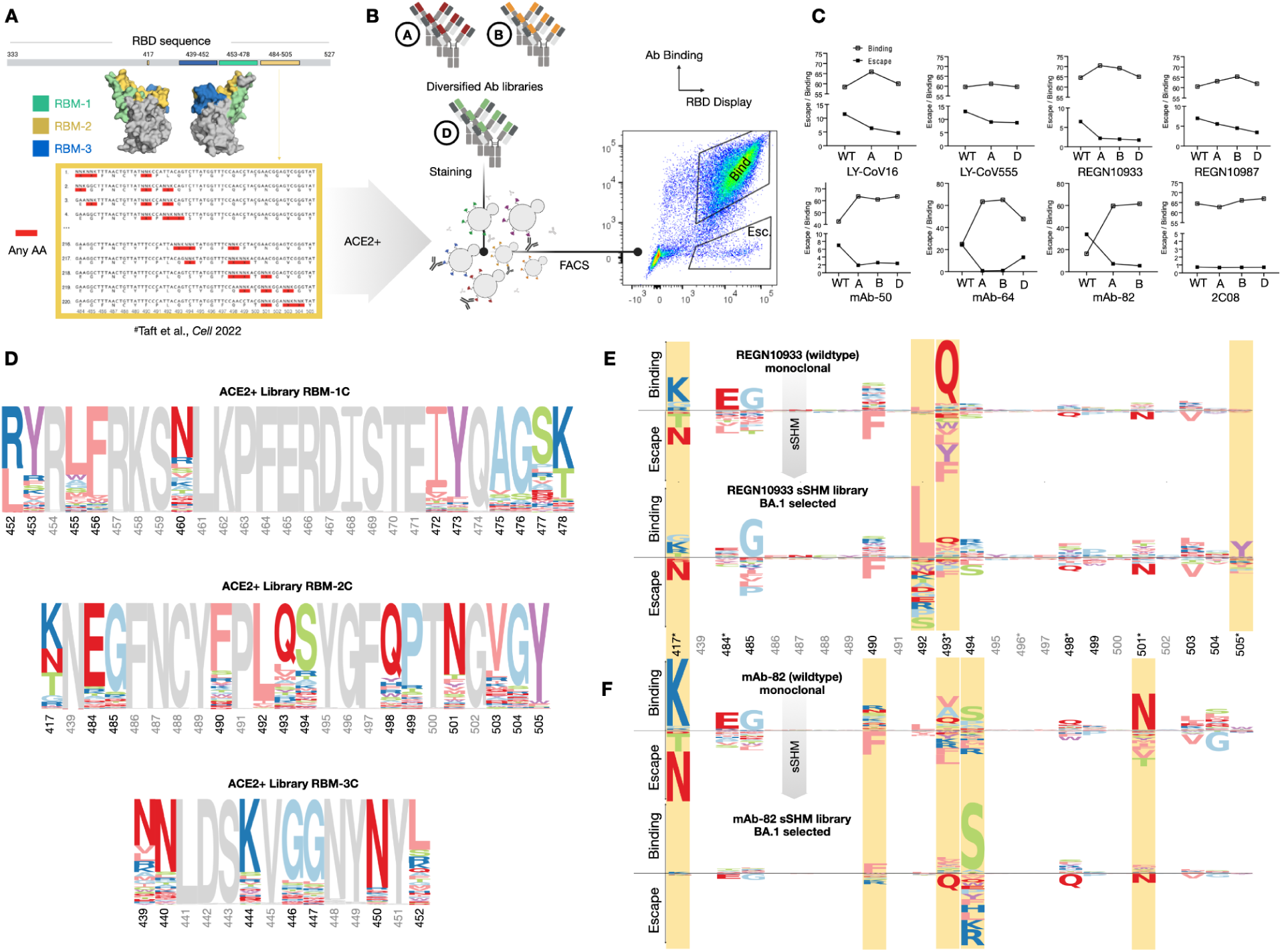
Coevolution of RBD variants and their mutational mapping to neutralizing antibodies. (**A**) Synthetic RBD variants are designed based on mutagenesis libraries targeting RBM-1, -2 and -3. (**B**) Antibody clonotype (sSHM) pools that were previously selected for binding to defined RBD variants (Alpha (A), Beta (B) and Delta (D)) are then screened by yeast surface display of synthetic RBD variants (pre-selected for binding to ACE2). Representative FACS plot of RBD variants labeled with an antibody clonotype pool, including sorting gates for binding and escape fractions. (**C**) The fraction (%) of the synthetic RBD variants that either bind or escape the parental (WT) antibody or antibody clonotype pools previously selected for binding to defined RBD variants. (**D**) Protein sequence logos of the RBD variants that maintain binding to ACE2 from the RBM-1C (top), RBM-2C (middle) and RBM-3C (bottom) libraries. Gray residues indicate positions that were conserved in the RBM library design. (**E, F**) Enrichment and depletion analysis of specific RBD positions based on binding or escape to either parental (WT) or BA.1-selected clonotype pool of REGN10933 (E) or mAb-82 (F). Gray position numbers correspond to positions not mutated in the RBM library, while an asterisk indicates positions relevant to BA.1

A combination of yeast RBD mutagenesis libraries were selected for binding or non-binding to soluble ACE2 receptor by FACS; the ACE2-binding population of RBD variants were subsequently screened with selected monoclonal antibodies or antibody clonotype pools for binding and escape (non-binding) by FACS (**Fig. 3B**, **Fig. S6**). When comparing the monoclonal precursors with their selected clonotype libraries, the diversified antibody libraries either maintained or exceeded the fraction of bound RBD variants, while reducing or maintaining escape as measured by flow cytometry (**Fig. 3C**). Targeted deep sequencing of the RBD libraries that maintain binding to ACE2 reveal the evolvability of some positions and residues (**Fig. 3D**). By normalizing for the distribution of the ACE2-binding RBD fraction (background normalization), we observe specific mutations that drive binding or escape to a given antibody or antibody clonotype pool (**Fig. 3E, F**).

As expected, we identified changes in the binding landscape particularly in positions that were selected for during the process of sSHM (for Beta or BA.1), such as mutations in RBD positions: 417(N), 484(K/A), 493(R/Q), 498(R/Q), 501(Y) and 505(H/Y) (**Fig. 3E, F**). Compensatory changes observed outside of these positions, such as in 485, 490, 492, 499, 503, 504 are therefore also of interest for deciphering potential coevolutionary adaptations. For instance, applying sSHM to REGN10933 caused a shift in the binding landscape with particular importance of RBD position 492, in which any mutation appears in this position to drive escape (**Fig. 3E**), while previous studies employing deep mutational scanning (DMS) did not show 492 being associated with escape to REGN10933 (*18*). We also observed a decrease in escape from RBD position 417, albeit to a smaller degree than expected, with K417N fractions appearing diminished, while K417T is not detected after sSHM. Similarly, RBD positions 484 and 493 appear to have lower relevance, while positions 494 and 505 show a higher degree of escape mutations (**Fig. 3E**). Applying sSHM to mAb-82 also substantially altered its binding landscape, as it appears to no longer be susceptible to the RBD escape mutations of K417N/T and has a strong reduction in reliance on positions 493 and 501 (**Fig. 3F**), while exposing it to changes in position 494. Positions 493, 494 and 452 are in close proximity within the structure of the RBD (**Fig. S7**), and as shown earlier for REGN10933 (**Fig. 2E**), mAb-82 exhibits an increased susceptibility to changes in RBD L452, hinting towards a similar origin of these changes.

### Synthetic convergent evolution identifies RBD variants with broad antibody escape

Next, we sought to evaluate the hypothesis that specific mutations drive the escape within a large number of the tested antibodies and corresponding antibody libraries. Convergent evolution, the independent accumulation of common mutations across several unrelated and parallel SARS-CoV-2 lineages (*19*), has been observed in RBD positions such as N501 (*20*), E484 and K417 (*21*) and across a multitude of positions in Omicron sublineages (*22*). Convergent mutations are often associated with enhanced binding to ACE2 and/or escape from common neutralizing antibodies (*23*). We, therefore, sought to identify convergent (overlapping) RBD mutations that drive escape to the polyclonal antibody pools obtained by sSHM. RBD variants were clustered based on RBM-1, -2, and -3 sequence similarity (Levenshtein edit distance), resulting in a highly branched network. Limiting the network to only include RBD sequences escaping from at least 10, 8 or 7 monoclonal antibodies or antibody clonotype pools, results in dense cluster formations (**Fig. 4A, B**). Selected clusters are visualized as sequence logos, the cluster identity is given by the number of shared mutations, or “driver” mutations, with additional positions of diversity potentially contributing to RBD escape, such as clusters containing and T478K, with limited diversity in Y453 and I472 for RBM-1 (**Fig. 4A, B left**). For RBM-2, mutations in 485, 492, 493 and 494 appear to drive multi-antibody escape, while K417 and E484 appear to remain mostly unmutated (**Fig. 4A, B, middle**). This in turn may be expected given that most antibody pools were selected for K417N and E484K adaptation, potentially weakening interactions with the wild-type residues K417 and E484. In RBM-3, all clusters exhibit diversity in position 452, either with or without the involvement of 439 and 444, 446 and 447, while positions 440 and 450 remain largely constant. However, most antibodies and antibody clonotype pools exhibited little escape from the RBM-3 library (**Fig. 4A, B, right**), with the exception of the RBM-3 class antibodies, REGN10987_(sSHM)_ and mAb-50_(sSHM)_. Similarly, the majority of antibodies escaped by the respective RBM libraries could be mapped back to their respective epitope class (i.e., class 1 antibodies are escaped by RBM-1 variants, class 2 antibodies are escaped by RBM-2 variants). Nevertheless, we identified several synthetic RBD variants that escape antibodies outside of their given epitope class (the epitope is defined as any amino acid within 4.5Å of the antibody when bound to the Spike protein (*4*)). The variants [Y453V, Y473L, A475E], [G485D, L492S, V503E] and [N439S, L452P] escaped antibodies LY-CoV555, REGN10987 and LY-CoV016, respectively (**Fig. 4C**), despite the fact that none of the individual (single) amino acid changes were previously observed as being associated with escape (*24*, *25*). Escape is therefore likely to result from cumulative, epistatic changes to the overall RBD structure, thus stressing the importance of probing the combinatorial sequence space to comprehensively map and predict prospective escape variants.

**Figure 4.**
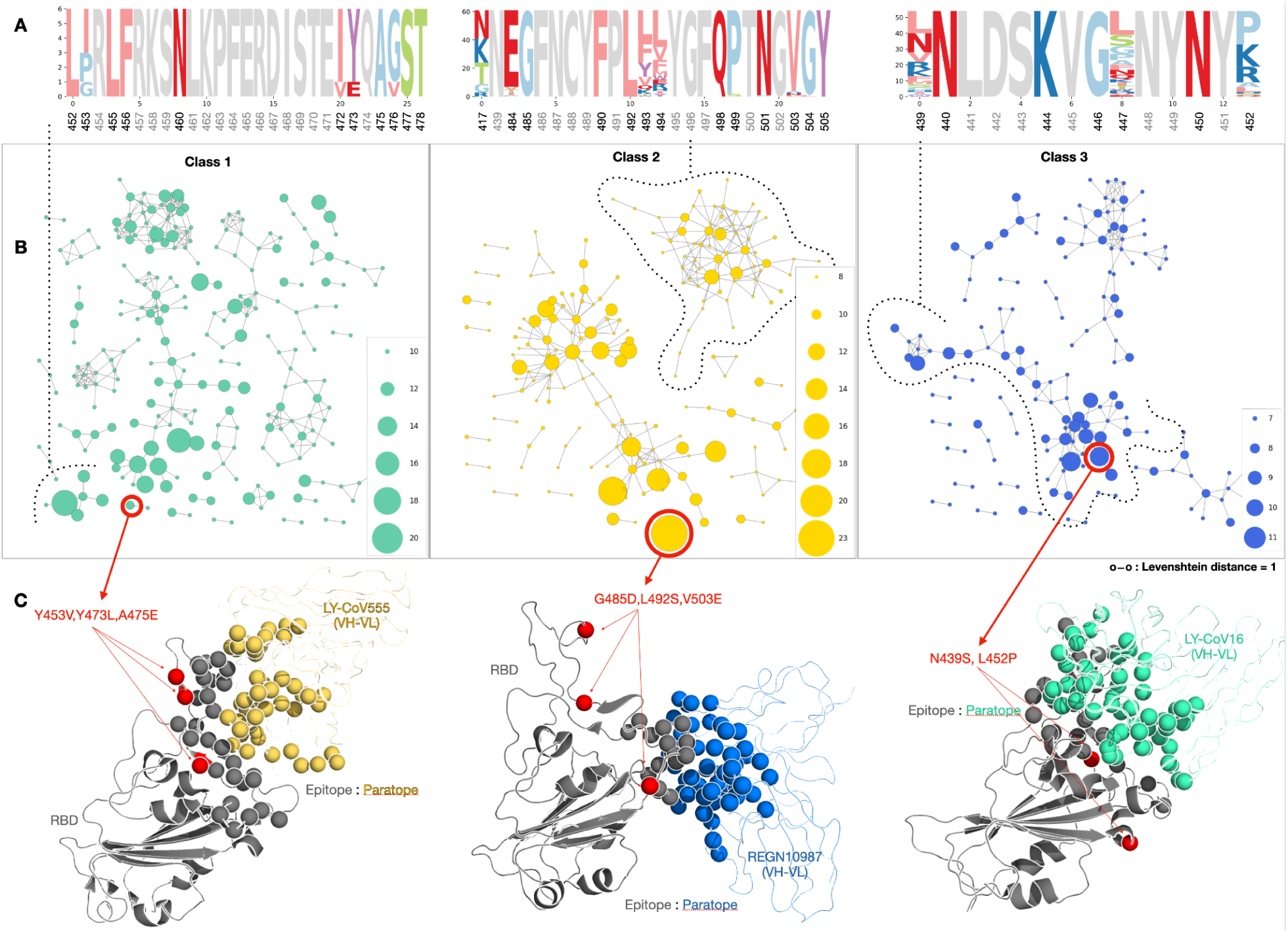
RBD variants with broad antibody escape profiles assemble into clusters. (**A**) Representative logo plots of a specific cluster (dotted line). (**B)** Each dot represents an RBD sequence that was found to escape in *n* datasets. (**C)** Representative crystal structure indicating an antibody’s epitope (yellow, blue or cyan spheres) and the RBD (RBM) class paratope (gray spheres). Red spheres indicate the position of the chosen set of mutations within the RBD structure.

### Deep learning predicts coevolved antibody escape profiles across a highly diverse landscape of RBD variants

Even with manageable RBD library sizes of ∼10^7^, oversampling requirements rapidly approach the limits of experimental yeast display screening. Therefore, to identify potential evolutionary trajectories, we trained deep learning models to make accurate predictions of antibody escape across a highly diverse landscape of potential RBD variants (>10^10^). To this end, we developed RBD-pLM, which combines a protein sequence language model with masked label modeling and inter-attention. Based on the sequence of an RBD variant, RBD-pLM is able to predict the binding or escape of 38 different antibody types (monoclonal antibodies or clonotype pools originating from this study and monoclonal antibodies from our previous study (*17*)) and ACE2 (**Fig. 5A**). The training, validation, and test datasets were derived from the deep sequencing of experimental screens (yeast display of RBD mutagenesis libraries for ACE2 / antibody binding or escape). The learning task was framed as a multi-label classification with partial labels. Multi-label classification is a generalization of multiclass classification that assigns multiple non-exclusive binary labels (1 or 0) to a given input. The model inputs were RBD sequences and the labels indicate binding (0) or escape (1) to antibodies and/or antibody pools and ACE2. Multi-label classification with partial labels is a more difficult task in which the full label set (38 antibodies + ACE2) lacks one or more antibody labels for each input (RBD sequence). The majority of RBD sequences are sparsely labeled, with label frequency per sequence exhibiting the long-tailed distribution typically associated with multi-label learning (**Fig. S8**). Furthermore, we observed uneven edit distance distribution and class balance across the different antibodies and RBM-1, -2 and -3 **(Fig. S9, S10**). We hypothesize that the RBD-pLM architecture might improve generalizability, overcoming limitations such as data scarcity or imbalance for certain antibody and RBD datasets.

**Figure 5.**
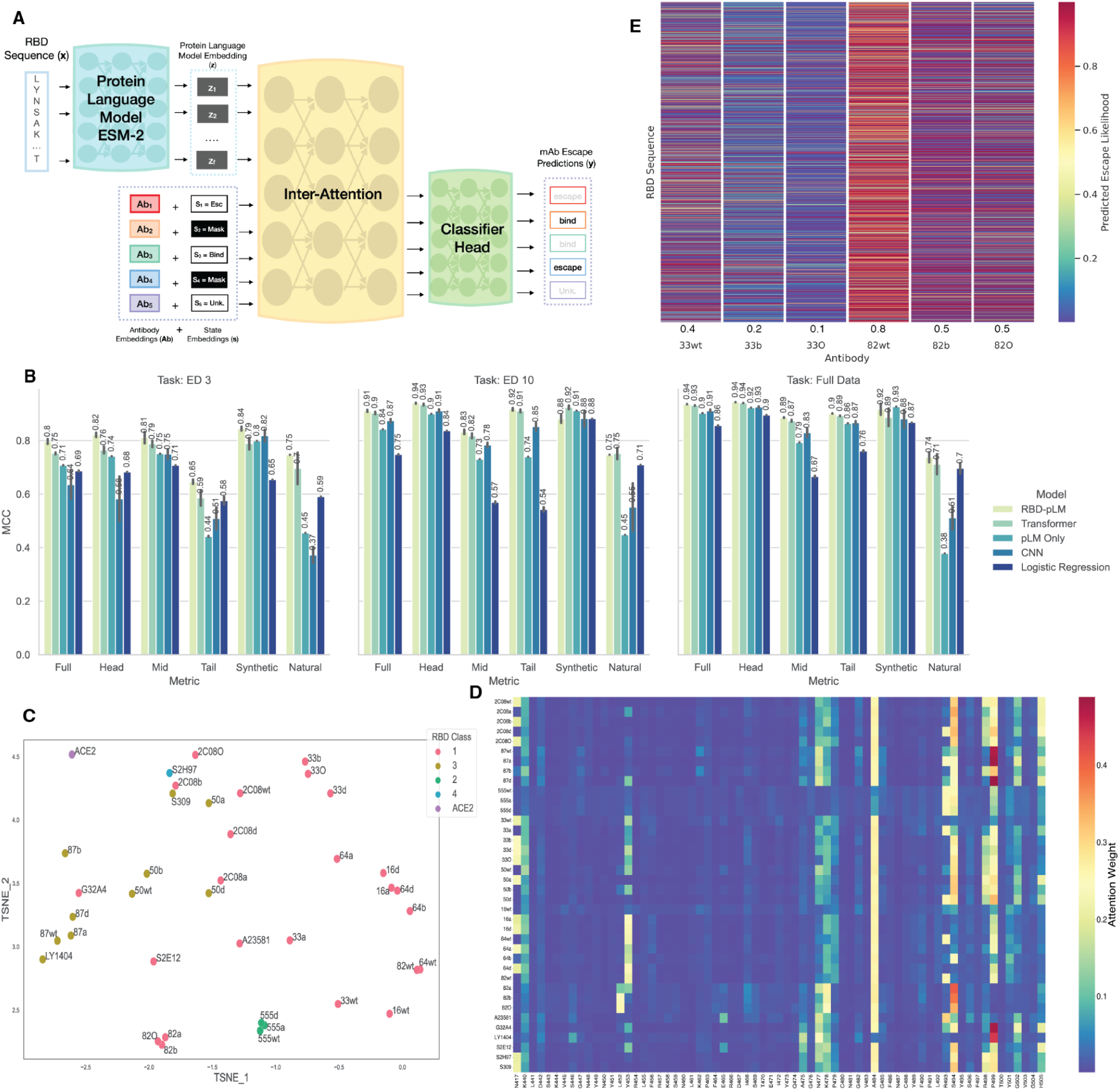
Deep learning accurately predicts antibody binding and escape of SARS-CoV-2 RBD variants. (**A**) Schematic diagram of RBD-pLM model. Model input consists of i) RBD variant sequences and ii) antibody/ACE2 state labels (binding, escape, or unknown). RBD variant sequences are fed to the ESM-2 (*26*) protein language model to obtain hidden representation **Z**. 85% of antibody state profiles are masked and the resulting vector is combined with learned antibody/ACE2 embeddings to produce **Ab**. **Ab** and **Z** are fed to the inter-attention mechanism, a combination of self- and cross-attention (**Fig S11**). The model is tasked with recovering the masked antibody/ACE2 binding states during training and predicting all labels during inference. **(B)** Machine learning tasks assessing the ability to generalize to a higher mutational load. For learning tasks ED 3 and ED 10, train and test sets were split via thresholding on Edit Distance (ED) from Wu-Hu-1 sequence. The training set consists of sequences with mutations at and below the threshold while the test set contains sequences with a higher number of mutations. Models were also trained with access to the full mutational range of the data (Full Data). Full Data train and test sets were split according to label. Predictive performance is evaluated by Matthew’s Correlation Coefficient (MCC). MCC is calculated for all antibodies and for the most (Head), medium (Mid), and least (Tail) frequent antibody labels in the training set. Performance is also assessed on external experimentally validated variants of Synthetic (*17*) and Natural (*31*) lineage. Reported metrics refer to mean test set performance of three separate training runs using different random seeds. Error bars correspond to 95 % confidence intervals. **(D)** Learned antibody embeddings were extracted from RBD-pLM and clustered using the t-distributed Stochastic Neighbor Embedding (t-SNE). Antibodies generally cluster by epitope class and occasionally by clonotype pools. **(E)** Heatmap of RBD-pLM predicted antibody escape probabilities for REGEN10933 and mAb-82 wild-type and sSHM antibodies. Shown are the escape probabilities for 259,907 synthetic Beta lineage RBD variants. The binding/escape prediction threshold is set to 0.5. Above 0.5 is considered escape (red). 0.5 or below is considered binding (yellow to blue). Synthetic RBD lineages are generated as previously described (*17*). A summary statistic describing the average number of escaping RBD variants is calculated per target (antibodies and ACE2) and reported at the bottom of the columns. **(F)** RBD-pLM attention weights for RBD variant BA1.1 identifies residues implicated in antibody binding and escape. Heatmap reflects the summed attention weight over all model layers for a given position in the sequence. Antibody abbreviations: 16: LY-CoV16, 33: REGN10933, 87: REGN10987, 555: LY-CoV555. Variant abbreviations: a: Alpha, b: Beta, d: Delta, O: Omicron. sSHM antibody nomenclature indicates the selection variant e.g. 16d refers to LY-CoV16 selected on Delta. WT refers to wild-type antibodies.

RBD-pLM consists of two main tracks, one encoding RBD sequences and the other encoding the label set (antibody or ACE2 binding or escape) via masked label modeling (**Fig 5A**). RBD protein sequences are encoded using the pretrained protein language model ESM-2, as its embeddings have been shown to contain rich structural, evolutionary and functional information (*26*). In parallel to the RBD representation, RBD-pLM encodes the label set in stages via label embedding, binding state embedding, and masked label modeling (*27*). First, the labels are treated as an additional 39 tokens, for which embeddings are learned. Next, binding state embeddings are learned for the label vector (bind, escape, or unknown). A subset of the binding state embeddings are then masked out (e.g. flipped to unknown) before being added to the label embeddings. The RBD sequence embeddings are then mixed with the label embeddings via inter-attention, which is a combination of self- and cross-attention (**Fig. S11**). Finally, similar to masked language modeling, RBD-pLM is tasked during training with recovering the masked label binding states. Other important aspects of RBD-pLM include an element-wise classification head and the ability to learn from partially labeled data, among others (**see Materials and Methods**). In addition to RBD-pLM, we trained and evaluated several baselines including the protein language model without antibody embeddings and inter-attention (pLM only), a standard transformer, a convolutional neural network (CNN), and logistic regression.

Mutational load can dynamically increase during viral evolution, as evidenced by the emergence of highly mutagenized variants such as BA.1 and other Omicron sublineages (e.g., BA.2, BA.4/5, XBB, BQ.1.1), as well as the recently reported BA.2.86 and JN.1 (*28*, *29*). It is therefore desirable for a machine learning model of viral coevolution to generalize to a larger mutational space than that of its training set. To interrogate generalizability, three learning tasks were curated. Two of the learning tasks implement an edit distance threshold to separate the train and test sets. Models were trained on RBD sequences with an edit distance at or below three (ED-3) or ten mutations (ED-10) to the ancestral Wu-Hu-1 sequence, and their performance was tested on mutations above the threshold, with the maximum number of mutations being 23 (**Fig. 5B**). The final learning task split the train and test sets based on the label set distribution, thereby training on the full mutational range of the data (full data). Predictive performance was primarily assessed based on Matthew’s Correlation Coefficient (MCC) values (*30*) for the test set, the most frequent (head), medium frequency (mid), and least frequent (tail) antibody labels. Frequency bins are calculated according to training set label occurrence. Additional metrics including F1, Hamming Loss, Jaccard index, frequency-binned and aggregate scores were also calculated (**Table S3**). Classification performance for each of the antibody and ACE2 targets were also determined (**Table S4**). RBD-pLM outperforms the baselines for the majority of tasks and metrics. RBD-pLM performance improvements over the baselines are noteworthy for antibodies with small and unbalanced data. This is apparent in the head, mid, and tail summary metrics (**Fig. 5B**) and particularly at the individual antibody level (**Table S4**). For example, RBD-pLM yields an MCC of 0.85 ± 0.01 for Delta-selected 2C08_sSHM_ on the ED-3 task, while the most competitive baseline, the transformer, yields 0.66 ± 0.02. Similar results are seen for numerous antibodies, including Beta-selected REGN10933_sSHM_ (RBD-pLM 0.51 ± 0.06 and transformer 0.34 ± 0.02) and Delta-selected mAb-64_sSHM_ (RBD-pLM 0.70 ± 0.04 and transformer 0.58 ± 0.04). Furthermore, test performance generally increased with an increasing edit distance threshold and the highest performance was observed for models trained on the full mutational range (**Fig. 5B**).

Models were further validated on two previously published experimental datasets consisting of synthetic (*17*) or natural RBD variants (*31*). The synthetic variants consist of 46 RBD sequences derived from Alpha, Beta, and Kappa lineages and were experimentally tested for binding or escape to four antibodies (LY-CoV16, LY-CoV555, REGN10933, and REGN10987) and ACE2 (*17*). The natural variants includes 11 RBD sequences from various Omicron lineages experimentally tested for binding or escape to five antibodies (LY-CoV1404, S309, LY-CoV555, REGN10933, REGN10987) and ACE2 (*31*). RBD-pLM trained with full data accurately predicted antibody escape and ACE2 binding for the synthetic (MCC_full_ 0.92 ± 0.02) and natural variants (MCC_full_ 0.74 ± 0.02) (**Fig. 5B**), the latter being particularly challenging since the training data consisted of mutational libraries designed prior to the emergence of Omicron BA.1 (*17*). Logistic regression performs surprisingly well on the natural variants (MCC_full_ 0.70 ± 0.02) when trained with a large dataset. A potential cause may be its focus on low level patterns, including the large number of mutations compared to ancestral Wu-Hu-1 RBD. In contrast, it is unable to perform well on the large test set, which contains a wide distribution of RBD mutations and nuanced sequence motifs.

Next, we used t-distributed Stochastic Neighbor Embedding (t-SNE) to cluster antibody and ACE2 embeddings learned by RBD-pLM, which revealed several trends (**Fig. 5C**). In most cases, wild-type antibodies cluster together by class, while the variant-selected antibody clonotype pools are often distant from their respective wild-types (e.g. mAb-82) and sometimes even cluster with similar clonotype pools (e.g. antibody clonotype pools of LY-CoV16_sSHM_ and mAb-64_sSHM_). For example Omicron BA.1-selected 2C08_sSHM_ clusters away from its wild-type counterpart and closer to antibodies that were originally discovered against Wu-Hu-1 and retain binding to Omicron BA.1, such as S2H97 and S309.

Attention heatmaps can be used to help interpret transformer-based deep learning architectures and have previously been shown to correlate with protein folding, target binding sites, and biophysical properties (*32*). Model attention weights, at a high level, can represent how correlated two amino acids may be with respect to structure and function. Using BA1.1 as an example, inspection of RBD-pLM cross-attention weights, between antibodies and RBD sequence amino acids, highlight residues largely implicated in ACE2 binding and antibody escape (**Fig. 5D**). For example, K417N and Q493R, are respectively present and absent in BA1.1, and are shown to escape Class 1 and 2 antibodies and S309 (Class 3) (*5*). N501Y contributes to ACE2 binding but is absent (*33*). Additionally, N450, L452, F490, and P499 are related to antibody escape, and are suspected to be major antigenic sites (*34*). RBD-pLM also places significant weight on positions implicated in sSHM-derived antibody binding and escape such as 501Y and 484A. Furthermore, a shift in escape profile attention is observed for antibody clonotype pools compared to wild-type antibodies (e.g. mAb-82).

We have previously demonstrated that synthetic lineages with predicted ACE2-binding can be generated and used to simulate plausible evolutionary trajectories of SARS-CoV-2 RBD (*17*, *35*). We used this approach to predict binding and escape of antibodies across various synthetic lineages (**Fig. S12**); in particular, we highlight the escape profiles of REGN10933 and mAb-82 and their respective clonotype pools selected for binding to Beta or Omicron BA.1 on Beta-derived lineages consisting of ∼260,000 synthetic RBD variants (**Fig. 5E**). The findings reveal a nuanced landscape of antibody-variant interactions. Notably, the sSHM variant-selected antibody pools generally demonstrated a lower escape rate compared to their wild-type counterparts, suggesting that antibody affinity maturation and antibody breadth are not evolutionarily mutually exclusive.

## DISCUSSION

We have established a synthetic coevolution system to interrogate the mutational landscape of human neutralizing antibodies and SARS-CoV-2 RBD. By utilizing protein mutagenesis, high-throughput screening and deep sequencing, we were able to generate large-scale datasets of antibody and RBD variants.

We deployed sSHM for antibody library generation and mammalian surface display screening to identify antibody clonotypes that bind to different RBD variants. Even for antibodies that had lost nearly all detectable binding to Omicron BA.1, we could recover clonotype variants with high affinity capable of neutralization. For example, just a single point mutation in the HCDR3 of REGN10933 (T102S) was sufficient to restore binding and neutralization to BA.1. This supports the fact that in human immune responses, individuals that have been previously vaccinated or infected possess memory B cells that can undergo somatic hypermutation to evolve antibodies with high affinity and neutralization upon exposure to new SARS-CoV-2 variants (*2*). Since a minimal amount of mutations may be sufficient to recover binding of a neutralizing antibody to a new SARS-CoV-2 variant, synthetic coevolution (re-engineering) could be performed on a previous clinically used antibody therapeutic (e.g., REGN10933) to produce an updated version (*36*). Such an approach would be analogous to updating seasonal vaccines (i.e., COVID-19 and influenza), as the re-engineered antibody may retain similar drug developability and safety profiles and offer a possible strategy for accelerated regulatory approval. An example of this was the monoclonal antibody AGD2 (precursor of ADG20/adintrevimab), which exhibited broad reactivity to pre-Omicron variants (*37*), but resulted in a substantial loss of neutralization potency to Omicron variants and was subsequently withdrawn from clinical development (*38*). A process very similar to sSHM was used to re-engineer ADG20 to have broader reactivity to Omicron variants, this resulted in the antibody VYD222 (pemivibart) (*39*) which recently received FDA approval (emergency use authorization) (*40*)

To perform coevolution, we used yeast display screening of RBD combinatorial mutagenesis libraries to map the mutational landscape of antibody escape. Wildtype precursor antibodies and their evolved sSHM-derived variants showed distinct RBD escape profiles, revealing different RBD positions that are most susceptible to escape. This further showcases the immune-evasive potential of SARS-CoV-2 to undergo adaptive and compensatory coevolution to escape newly evolved neutralizing antibodies, while still maintaining the capacity to infect host cells via binding to ACE2. This is in line with natural coevolution: selective pressure exerted by human population-level humoral immunity, which is dynamic and constantly evolving, is one of the primary driving forces behind the continued emergence of multiple and simultaneously occurring SARS-CoV-2 variants and lineages (e.g., Omicron sublineages) (*23*).

While experimental screening by yeast display generated large and diverse datasets of synthetic RBD variants and their associated antibody binding and escape profiles, this only represented a minor fraction of the theoretical protein sequence space of the RBD. With the continued emergence of new variants with a high mutational load, such as the recently reported BA.2.86 and JN.1 (*28*, *29*), it becomes increasingly important to interrogate much more of the possible mutational landscape of SARS-CoV-2. Therefore, we trained deep learning models that were capable of making predictions on antibody binding and escape and extrapolate much deeper into the RBD protein sequence space. However, this was complicated by inherent limitations of experimental screening, one of which is that for the majority of RBD variants it was only possible to recover binding or escape labels for few antibodies or antibody pools (i.e., five or fewer out of 38). Furthermore, RBD variant data was often highly imbalanced within a given classification group (antibody binding or escape). To address these difficulties, we leveraged a protein language model, masked label modeling and inter-attention to construct the RBD-pLM. Self-supervised pretrained protein language models have been shown to encode functional, structural, and evolutionary information of proteins and to produce meaningful embeddings useful for downstream tasks (*26*). Due to the ability to efficiently learn relationships between inputs, we employed masked label modeling and inter-attention to embed antibodies and ACE2, while simultaneously learning relationships between the antibodies themselves and between the antibodies and RBD sequences. Through several computational trials, we demonstrated the ability of RBD-pLM to extrapolate to a broader mutational sequence space and perform well on external experimental data sets. We additionally probed RBD-pLM using explainable artificial intelligence techniques that could recover biologically meaningful trends, including for example, model attention weights highlighting the importance of RBD positions K417N and Q493R for escape to multiple antibodies (*5*). Our work also demonstrates the effectiveness of transfer learning and careful model design to exploit relationships between multiple classification groups and efficiently learn from a highly imbalanced dataset. Using protein language models for data efficient protein engineering is an emerging area, including the use of parameter efficient fine-tuning to substantially reduce computing requirements for large language models (*41–43*).

## MATERIALS AND METHODS

### Mammalian cell culture

Generation of nPnP founder cell line has been previously described (*10*). Electroporation was performed with the 4D-Nucleofector System (Lonza) using SF Cell line 4D-Nucleofector X Kit L or X Kit S (Lonza, V4XC-2024, V4XC-2032) with the program CQ-104. Previous to nucleofection, cells were centrifuged at 150xg for 5 minutes and washed once with pre-warmed (37 °C) Opti-MEM I Reduced Serum Medium (Thermo, 31984-062) at 1 mL per 1x10^6^ cells. The resulting cell pellet was then resuspended in SF buffer according to the manufacturer’s description, after which Alt-R gRNA (sequence: ATATGACTCCTTCGACTCGA) and (3.5-4.5 pmols per 1x10^6^ cells) HDR donors were added. Cells were analyzed for HDR events approx. 3-4 days post transfection.

Expi293 culture and transfections were carried out according to the manufacturer’s specifications (ThermoFisher, Cat#A14525 and as described previously (*44*)).

### Cloning and assembly of antibody clonotype (sSHM) libraries

Precursor HDR plasmids were cloned as follows: Truncated versions, missing the sequence between HCDR2 and HCDR3 were synthesized and ligated into PCR amplified pHDR backbone by Gibson cloning. Oligopools containing the diversified HCDR2 and HCDR3, respectively, were mixed at isomolar ratios of 10 pmols each. After 5 PCR cycles using Q5 polymerase (NEB #M0491) with 5X GC Buffer with a melting temperature of 72°C, PCR products were purified by silica spin columns. Plasmids containing truncated antibody sequences were resuspended to 75 ng/µL each. Golden Gate Assembly using PaqCI (NEB #R0745) and T4 Ligase (NEB #M0202) was performed as recommended by the manufacturer (using recommendations for “simple-moderate assembly difficulty”). Ligated plasmids were desalted over MilliQ deionized water for 1 hour prior to transformation into fresh electrocompetent cells (DH10Beta ElectroMAX).

### Screening mammalian display antibody libraries for binding to RBD variants

After an incubation time of 3 days, FACS (BD FACSMelody Cell Sorter) was performed to isolate PnP cells for antibody surface expression and display. Staining was performed with 2x10^5^ cells each and 1:100 anti-hIgG-AF488 (Jackson ImmunoResearch, 109-545-003) for detection of antibody display. After recovering cells for at least 3 days, FACS (BD FACSAria Fusion Cell Sorter / BD FACSMelody / Sony MA900) was performed on PnP cells expressing surface antibody and binding to soluble RBD protein. Cells were stained with 100 nM biotinylated RBD antigen, 1:200 Strepavidin-AF647 / 1:200 Streptavidin-PE and 1:100 anti-hIgG-AF488. Sorting for RBD binding was repeated at least once to obtain a more pure population.

### Cloning and expression of RBD mutagenesis libraries for yeast surface display

Tiling mutagenesis libraries of the RBD (library RBM-1T, RBM-2T and RBM-3T) were obtained from our previous study (*17*). For libraries RBM-2_K417N/T_ and RBM-1_L452R+T478K_, pools of single-stranded oligonucleotides (ssODN) were designed, where each member of the pool contains one combination of the three ‘NNK’ codons. Additionally, a primer was used to place a K417N or T degeneracy by introducing an “AMN” codon for library RBM-2_K417N/T_. For library RBM-1_L452R+T478K_, positions L452R and T478K were fixed to represent the Delta lineage. ssODN pools were amplified by PCR. Insert and EcoRI-linearized plasmids (*17*) were concentrated and purified by silica spin columns (Zymo D4013), with repeated washing. The libraries were cloned and expressed in yeast by in vivo homologous recombination, as previously described (*45*) , using 4 μg each of plasmid and 8 µg insert DNA per 300 μl of electrocompetent EBY100 cells in a 2 mm electroporation cuvette.

### Screening yeast display RBD libraries for binding to ACE2

The diversity of library RBM-2_K417N/T_and RBM-1_L452R+T478K_ were approximately 1*10^6^ and 2.5*10^6^, respectively. Yeast cell surface expression of RBD was induced by growth in SG-UT medium at 23°C for 16-40 hours, as previously described (*45*). Approximately 2*10^8^ library cells were washed once with 1 mL wash buffer (Dulbecco’s PBS+ 0.5% BSA + 0.1% Tween20 + 2 mM EDTA) by centrifugation at 8000 x g for 30 s. Washed cells were stained with 50 nM biotinylated human ACE2 (Acros AC2-H82E6) for 30 minutes at 4 °C, followed by an additional wash. Cells were then stained with 2.5 ng/μl streptavidin-AlexaFluor 647 (Biolegend 405237) and 1 ng/μl anti-FLAG-PE (Biolegend 637310) for 30 minutes at 4 °C. Cells were subsequently pelleted by centrifugation at 8000 x g for 30s and kept on ice until sorting. Binding (ACE2+/FLAG+) and non-binding (ACE2-/FLAG+) yeast cells were sorted by FACS (BD FACSAria Fusion). Collected cells were immediately checked for purity and resorted if necessary.

### Screening yeast display RBD libraries for binding and escape to antibodies

RBD libraries pre-sorted for ACE2-binding were cultured and induced, as described above. Induced cells were washed once with DPBS wash buffer, followed by a 30 min incubation with 100 nM monoclonal antibody, or 100 nM of polyclonal antibodies (sSHM-derived clonotypes) at 4°C while shaking at 750 rpm. Following an additional wash, cells were resuspended in 5 ng/μl anti-human IgG-AlexaFluor647 (Jackson Immunoresearch 109-605-098) and incubated for 30 min at 4°C shaking at 750 rpm. Yeast cells were washed once more and resuspended in 1 ng/μl anti-FLAG-PE for another 30 min of incubation at 4°C. Cells were then pelleted by centrifugation at 8000 x g for 30s and kept on ice until sorting. Yeast cells expressing RBD that maintained antibody-binding (IgG+/FLAG+) or showed a complete loss of antibody binding (escape) (IgG-/FLAG+) were sorted by FACS (BD Aria Fusion or BD Influx). Collected cells were cultured in SD-UT medium for 16-48 hours at 30°C. Induction and sorting was repeated for multiple rounds until the desired populations of RBD variants showed purity for binding and escape (non-binding) to antibodies.

### Monoclonal antibody production and purification

Heavy chain and light chain inserts for REGN10933, REGN10987 (PDB: 6XDG) and LY-CoV16 (PDB: 7C01), LY-CoV555 (PDB: 7KMG), mAb-50, mAb-64, mAb-82 and 2C08 were cloned into pTwist transient expression vectors by Gibson Assembly. 30 mL cultures of Expi293 cells (Thermo, A14635) were transfected according to the manufacturer’s instructions. After 5-7 days, dense Expi293 cultures were centrifuged at 300 x g for 5 minutes to pellet the cells. Supernatant was filtered using Steriflip 0.22 µm (Merck, SCGP00525) filter units. Using protein G purification, Expi supernatant was directly loaded onto Protein G Agarose (Pierce, Cat# 20399) gravity columns, washed twice with PBS and eluted using Protein G Elution Buffer (Pierce, Cat# 21004). The eluted fractions were immediately neutralized with 1M TRIS-Buffer (pH 8) to physiological pH and quantified by Nanodrop 2000c for A280 nm absorption. Protein containing fractions were pooled and buffer exchanged using SnakeSkin dialysis tubing (10 MWCO, Pierce Cat#68100) followed by further dialysis and concentration using Amicon Ultra-4 10kDa centrifugal units (Merck, Cat# UFC801096), as described previously (*44*).

### Polyclonal antibody production and purification

Selected antibody clonotype (sSHM) libraries expressed in PnP cells were cultured as described previously in T-225 flasks containing 50 mL medium until maximum density (∼3*10^6^ cells/mL, >90% viability). Supernatant was centrifuged at 300g for 5 min, and then filtered using a SteriFlip. Using Protein G binding buffer, pH was adjusted to pH 5-6 to enable efficient binding of antibodies to the Protein G resin. Adjusted supernatant and Protein G were incubated at 4°C overnight while spinning in a rotator and then transferred to gravity columns. The remaining process was carried out as described above(see “Monoclonal antibody production and purification”).

### Pseudotyped virus neutralization assays

Neutralization assays were performed as previously described (*46*). Briefly, pseudotyped lentiviruses displaying respective spike proteins (codon-optimized and with 19aa C-terminal truncations) and packaging a firefly luciferase reporter gene were generated by the co-transfection of HEK293T cells using polyethylenimine. Media was changed 12-16 hours after transfection, and pseudotyped viruses were harvested from the supernatant at 48- and 72-h post transfection. These were then clarified by centrifugation and stored at -80C until use. Pseudotyped viruses titrated to produce approximately 100,000 relative light units (RLUs) were incubated with serial 3-fold dilutions of monoclonal antibodies for 60 min at 37C in a 96-well black-walled plate, and then 10,000 HEK293T-ACE2 cells were added to each well. Plates were then incubated at 37C for 48 h, and luminescence was measured using Bright-Glo substrate and lysis buffer (Promega) per the manufacturer’s protocol, on a GM-2000 luminometer (Promega).

### Deep sequencing of RBD libraries

Plasmid DNA encoding the RBD variants was isolated from yeast cells following the manufacturer’s instructions (Zymo D2004). Mutagenized regions of the RBD were amplified using custom oligonucleotides. Illumina Nextera barcode sequences were added in a second PCR amplification step, allowing for multiplexed high-throughput sequencing runs. Populations were pooled at the desired ratios and sequenced using Illumina 2 x 250 PE protocols (MiSeq or NovaSeq instruments).

### Targeted deep sequencing of antibody libraries

Targeted amplification of V_H_ libraries from PnP cells for deep sequencing were prepared as described previously (*47*). Briefly, genomic DNA was extracted from 1 x 10^6^ cells using a PureLink™ Genomic DNA mini Kit (Thermo, Cat: K182001). Extracted DNA was PCR amplified with a forward primer binding to the end of the leader sequence (PCR1 Fwd primer with Illumina adapter overhang) and a reverse primer annealing to the intronic region directly 3’ of the J segment (PCR1 Rev primer with Illumina adapter overhang). PCRs were performed with Q5 High-fidelity DNA polymerase (NEB, M0491L) plus GC buffer with the following reaction conditions: 1 cycle of 98°C for 30 sec; 20 cycles of 98°C, 60°C for 20 sec, 72°C for 30 sec; and a final extension at 72°C for 1 min; followed by 10°C for temporary storage. Products were checked on an agarose gel and subjected to a PCR-clean up step (DNA Clean and Concentrator-5, Zymo, D4013). A second PCR (PCR2) was performed to add Illumina index adaptors (Nextera UD Index kits (plate C)). Final PCR products were gel purified (2% TAE).

PCR1 Fwd primer with Illumina adapter overhang:

5’ **CCCTCCTTTAATTCCC**TGATTCTTTTTGTCTAAAGGTATCCTGTC 3’

PCR1 Rev primer with Illumina adapter overhang:

5’ **GAGGAGAGAGAGAGAG**GTTAAAAATAAAGACCTGGAGAGGCC 3’

PCR2 was performed using Nextera’s UD Index kits (plate C). **Bold text** indicates Illumina adapter sequences.

### Processing of antibody and RBD deep sequencing data, statistical analysis and plots

Sequencing reads were paired, quality trimmed and assembled using Geneious and BBDuk, with a quality threshold of qphred ≥ 25. Mutagenized regions of interest were then extracted using custom Python scripts, followed by translation to amino acid sequences.

The RBD sequences obtained from each of the libraries were pre-processed separately before being combined into the final training set used for model training and evaluation.

Statistical analysis was performed using R 4.0.1 (*48*) and Python 3.8.5 (*49*). Graphics were generated using the ggplot2 3.3.3 (*50*), matplotlib 3.6.3 (*51*), ComplexHeatmap 2.4.3 (*52*), pheatmap 1.0.12 (*53*), igraph 1.2.6 (*54*), RCy3 2.8.1 (*55*), stringr 1.4.0 (*56*), dplyr 1.0.6 (*57*), and RColorBrewer 1.1-2 (*58*) R package.

### Preprocessing RBD deep sequencing data for machine learning

Following deep sequencing and processing with BBDUK and python,, all RBD sequencing data was preprocessed with Python (3.8.5) (*49*) using pandas (1.5.3) (*59*) and numpy (1.23.1) (*60*). RBD variant sequences present in both binding and non-binding (escape) populations for a given antibody were excluded from subsequent machine learning steps. Two types of train/test split approaches were investigated: splitting via edit distance cutoff and splitting via label distribution. The labeled RBD sequencing data from this study was combined with labeled RBD sequencing data generated from our previous study (*17*).

#### Train test split via edit distance

Levenshtein distance, or edit distance (ED) from the ancestral Wu-Hu-1 RBD sequence was calculated for all sequences. An ED cutoff was used to separate the training and validation sets from the test sets. Models were trained on RBD sequences with less than or equal to the ED threshold number of mutations and tested on sequences with more mutations than the threshold. Threshold splits of ED-3 and ED-10 from the ancestral Wu-Hu-1 RBD sequence were evaluated. Test sets contain sequences with up to 23 mutations from the ancestral Wu-Hu-1 RBD sequence **(Fig. S9)**. Data set sizes are reported (**Table S2**)

#### Train test split via label distribution

A previously described train test split strategy was used that takes into account the label set distribution (*61*). This split strategy enables models to access the full mutational range of the data (full data) during training. The python package iterative-stratification (0.1.6) was used to separate the data into training, validation, and testing. Data set sizes are reported (**Table S2**).

### Machine learning model development and implementation

#### Multi-label classification with partial labels

We sought to establish a machine learning model capable of predicting binding or escape (non-binding) labels for RBD variant sequences for 39 targets: 38 antibodies (monoclonal antibodies or antibody clonotype pools) and ACE2 receptor. Therefore, the learning problem was framed as a multi-label classification task with partial labels. Multi-label classification is a generalization of multiclass classification that assigns multiple nonexclusive binary labels (1 or 0) to a given input. Partially labeled refers to RBD sequences for which only a subset of the 39 binding/escape labels are experimentally available. Missing and masked labels are marked as unknown (labeled as -1). Missing labels were excluded from the loss calculation during stochastic gradient descent.

#### Protein language model

The protein language model ESM-2 was used as the RBD sequence feature extractor as its embeddings have been shown to contain rich structural, evolutionary, and functional information (*26*). The pretrained 8 million parameter version of ESM-2 was selected due to validation set performance. All ESM-2 versions were tested, however, these provided only small increases in performance while requiring significantly more time and resources to train. The 201 amino acid RBD sequences were fed to the tokenizer, which appends additional tokens for the beginning and end of the sequence, respectively. After extracting the final hidden layer representation, the beginning and end of sequence tokens were removed resulting in a 201 by 320 matrix, where 201 is the RBD sequence length and 320 the ESM-2 embedding dimension. RBD sequence embeddings were then fed to the inter-attention module (described below).

#### Label embeddings and binding state embeddings

The 39 classification targets (38 antibodies/antibody clonotype pools and ACE2) are represented as a constant vector of length 39, in which integer elements range from 0 to 38. The constant label vector is encoded with a learned embedding layer (referred to as **Ab**, see **Fig. 5)**. In parallel, the vector of binding state labels with 39 elements (one for each target) containing binding (0), escape (1), or unknown (-1) labels is masked via masked label modeling (discussed below) and encoded with a learned embedding layer. The label embeddings and (masked) binding state embeddings are then summed and fed to the inter-attention module (see below).

#### Masked label modeling

Masked label modeling closely resembles masked language modeling in which a fraction of the input tokens are masked out and the model is tasked with recovering the masked tokens during training (*27*). In masked label modeling, the full label vector (binding/escape state labels for 39 targets) is masked (set to -1) and a subset of known labels is probabilistically revealed. The masked label vector is encoded via learned embeddings (described above) and fed to the inter-attention module as a 39 token sequence matrix of size 39 by 320, where 320 is the embedding dimension. During training, the model is tasked with recovering the masked binding state identities. Previous works require a fully labeled training set, however we establish an implementation capable of learning with partially labeled data by ensuring unmasking and training loss calculations are only executed on the subset of known labels.

Three hyperparameters were tuned to control masked label modeling-based validation set performance. First, the initiation threshold **t**, ensures that only 25% of training examples will have a subset of their known labels revealed while the remaining 75 % of training examples will contain a fully masked label set. The second hyperparameter **q** (set to 3 in our experiments), refers to the quantity of known labels a training example must contain for a subset of the known labels to be revealed. Setting **q** above 2 forces the model to learn label co-occurrence relationships while preventing the trivial case that provides the model with the only known label in the label vector. The final hyperparameter **f** refers to the fraction of labels to reveal. If the conditions set by **t** and **q** are met, **f** percent of the known label set is revealed. In our experiments, we set **f** to 15 %.

#### Inter-attention mechanism

The inter-attention mechanism combines self-attention and cross-attention to evolve the embeddings of the RBD sequence and the labels in a parallel fashion (**Fig. S12**). A single inter-attention block consists of two tracks: one for the RBD sequence embedding and one for the target label embeddings. To promote stable training, at the beginning of each inter-attention block the RBD and label embeddings are concatenated and normalized via Layer Normalization (*62*). The embeddings are then separated and each embedding is subjected first to a self-attention layer and then a cross-attention layer in which it attends to the other embedding. For the learning tasks ED-10 and full data, RBD-pLM makes use of 3 inter-attention blocks in series. 2 inter-attention blocks are used for the ED-3 task.

Following the last inter-attention block, the RBD sequence embeddings and label embeddings are concatenated and subjected to a final self-attention layer. The label embeddings are then separated and fed to the classifier head.

#### Classifier head

We make use of an elementwise classifier head (*63*) that takes as input the label embeddings from the final inter-attention layer and outputs logits for each label. In-line with multi-label learning, Binary Cross Entropy is used as the loss function. Training loss is only computed for the subset of known labels that were masked out via masked label training.

#### Training

Models were coded and trained in Python (3.8.5) (*49*) using Pytorch (2.1.2) (*64*) with CUDA (11.8) (*65*). AdamW was used as the optimizer. Models were trained using the ETH Zurich Euler cluster. Training runs were performed in a distributed data parallel manner with a single node containing 4 GPUs. Models were trained in triplicates using different random seeds.

#### Baselines

Four baselines were evaluated: pLM Only, Transformer, CNN, and Logistic Regression. The pLM Only baseline encodes RBD sequences with the protein language model ESM-2 followed by a simple linear layer as its classifier head. This baseline serves to probe the contribution of the pLM. The Transformer baseline leverages attention, but does not make use of a pLM or masked label modeling. To mirror RBD-pLM, the embedding dimension is set to 320. The self-attention layers are followed by a linear layer of dimension 1024 and a classifier head. The CNN baseline is a convolutional neural network (*66*) with a 1D convolutional layer, with stride of 2 and a kernel of 3, followed by a linear layer of dimension 1024 and a classifier head. The Logistic Regression baseline serves as the simple baseline.

#### Performance metrics

Performance metrics including Matthew’s Correlation Coefficient (MCC), F1 score, Jaccard Index, and Hamming loss were calculated in aggregate for the full label set, for individual labels, and for subsets of the label set based on the most (Head), medium (Mid), and least (Tail) frequent labels (**Fig. 5, Table S3, Table S4**). MCC values range from -1 to 1. MCC was selected as a main performance metric due to its appropriateness for balanced and unbalanced classification tasks (*30*).

## ACKNOWLEDGMENTS

Computing resources and support from ETH Zurich (Euler cluster) are gratefully acknowledged. We would also like to gratefully acknowledge the ETH Zurich D-BSSE Single-Cell Facility and Genomics Facility, including Dr. Mariangela Di Tacchio, Renan Antonialli, Dr. Christian Beisel, Elodie Burcklen, Ina Nissen, and Mirjam Feldkamp. Some figures created with BioRender.com.

## Funding

This project is supported by Basel Research Centre for Child Health Fast Track Call for Covid-19 (to S.T.R.), Gemeinde Unterägeri via the ETH Zürich Foundation (to S.T.R), Swiss National Science Foundation (project: 310030_197941) (to S.T.R), and European Research Grant CoroNAb (to S.T.R. and B.M.)

## Author contributions

Conceptualization, R.E., M.M., and S.T.R.; investigation, R.E., M.M., M.D.O., D.J.S, J.H., B.G., J.M.T., M.P., C.R.W, T.B.; software, M.M., M.P., T.B.; supervision, B.M., S.T.R.; funding acquisition, S.T.R. B.M.; writing- original draft R.E., M.M. and S.T.R. Writing - review and editing: all authors.

## Competing Interests

S.T.R. may hold shares of Alloy Therapeutics and Engimmune Therapeutics. S.T.R. is on the scientific advisory board of Alloy Therapeutics and Engimmune Therapeutics. R.E., and B.G., and J.M.T. are employees of Enginmmune Therapeutics. C.R.W. is an employee of Alloy Therapeutics.

## Data and materials availability

The main data supporting the results in this study are available within the paper and its supplemental information. Raw deep sequencing data will be deposited to public databases (e.g. NCBI, Mendeley) upon manuscript acceptance. The code for this study is publicly available at https://github.com/LSSI-ETH/synthetic-coevolution.

Additional data files that support the findings of this study are available from the corresponding author upon reasonable request.

## Notes

https://github.com/LSSI-ETH/synthetic-coevolution

## REFERENCES

1. W. Ma, H. Fu, F. Jian, Y. Cao, M. Li, Immune evasion and ACE2 binding affinity contribute to SARS-CoV-2 evolution. Nat Ecol Evol, doi: 10.1038/s41559-023-02123-8 (2023).

2. C. I. Kaku, T. N. Starr, P. Zhou, H. L. Dugan, P. Khalifé, G. Song, E. R. Champney, D. W. Mielcarz, J. C. Geoghegan, D. R. Burton, R. Andrabi, J. D. Bloom, L. M. Walker, Evolution of antibody immunity following Omicron BA.1 breakthrough infection. Nat. Commun. 14, 2751 (2023).

3. M. Korenkov, M. Zehner, H. Cohen-Dvashi, A. Borenstein-Katz, L. Kottege, H. Janicki, K. Vanshylla, T. Weber, H. Gruell, M. Koch, R. Diskin, C. Kreer, F. Klein, Somatic hypermutation introduces bystander mutations that prepare SARS-CoV-2 antibodies for emerging variants. Immunity, doi: 10.1016/j.immuni.2023.11.004 (2023).

4. M. Cox, T. P. Peacock, W. T. Harvey, J. Hughes, D. W. Wright, COVID-19 Genomics UK (COG-UK) Consortium, B. J. Willett, E. Thomson, R. K. Gupta, S. J. Peacock, D. L. Robertson, A. M. Carabelli, SARS-CoV-2 variant evasion of monoclonal antibodies based on in vitro studies. Nat. Rev. Microbiol., doi: 10.1038/s41579-022-00809-7 (2022).

5. Y. Cao, J. Wang, F. Jian, T. Xiao, W. Song, A. Yisimayi, W. Huang, Q. Li, P. Wang, R. An, J. Wang, Y. Wang, X. Niu, S. Yang, H. Liang, H. Sun, T. Li, Y. Yu, Q. Cui, S. Liu, X. Yang, S. Du, Z. Zhang, X. Hao, F. Shao, R. Jin, X. Wang, J. Xiao, Y. Wang, X. S. Xie, Omicron escapes the majority of existing SARS-CoV-2 neutralizing antibodies. Nature 602, 657–663 (2022).

6. M. Ragonnet-Cronin, R. Nutalai, J. Huo, A. Dijokaite-Guraliuc, R. Das, A. Tuekprakhon, P. Supasa, C. Liu, M. Selvaraj, N. Groves, H. Hartman, N. Ellaby, J. Mark Sutton, M. W. Bahar, D. Zhou, E. Fry, J. Ren, C. Brown, P. Klenerman, S. J. Dunachie, J. Mongkolsapaya, S. Hopkins, M. Chand, D. I. Stuart, G. R. Screaton, S. Rokadiya, Generation of SARS-CoV-2 escape mutations by monoclonal antibody therapy. Nat. Commun. 14, 3334 (2023).

7. V. Van, A NEW EVOLUTIONARY LAW. (1973).

8. C. O. Barnes, C. A. Jette, M. E. Abernathy, K.-M. A. Dam, S. R. Esswein, H. B. Gristick, A. G. Malyutin, N. G. Sharaf, K. E. Huey-Tubman, Y. E. Lee, D. F. Robbiani, M. C. Nussenzweig, A. P. West Jr, P. J. Bjorkman, SARS-CoV-2 neutralizing antibody structures inform therapeutic strategies. Nature 588, 682–687 (2020).

9. A. J. Schmitz, J. S. Turner, Z. Liu, J. Q. Zhou, I. D. Aziati, R. E. Chen, A. Joshi, T. L. Bricker, T. L. Darling, D. C. Adelsberg, C. G. Altomare, W. B. Alsoussi, J. B. Case, L. A. VanBlargan, T. Lei, M. Thapa, F. Amanat, T. Jeevan, T. Fabrizio, J. A. O’Halloran, P.-Y. Shi, R. M. Presti, R. J. Webby, F. Krammer, S. P. J. Whelan, G. Bajic, M. S. Diamond, A. C. M. Boon, A. H. Ellebedy, A vaccine-induced public antibody protects against SARS-CoV-2 and emerging variants. Immunity 54, 2159–2166.e6 (2021).

10. R. A. Ehling, C. R. Weber, D. M. Mason, S. Friedensohn, B. Wagner, F. Bieberich, E. Kapetanovic, R. Vazquez-Lombardi, R. B. Di Roberto, K.-L. Hong, C. Wagner, M. Pataia, M. D. Overath, D. J. Sheward, B. Murrell, A. Yermanos, A. P. Cuny, M. Savic, F. Rudolf, S. T. Reddy, SARS-CoV-2 reactive and neutralizing antibodies discovered by single-cell sequencing of plasma cells and mammalian display. Cell Rep. 38, 110242 (2022).

11. M. Pogson, C. Parola, W. J. Kelton, P. Heuberger, S. T. Reddy, Immunogenomic engineering of a plug-and-(dis)play hybridoma platform. Nat. Commun. 7, 12535 (2016).

12. D. Planas, D. Veyer, A. Baidaliuk, I. Staropoli, F. Guivel-Benhassine, M. M. Rajah, C. Planchais, F. Porrot, N. Robillard, J. Puech, M. Prot, F. Gallais, P. Gantner, A. Velay, J. Le Guen, N. Kassis-Chikhani, D. Edriss, L. Belec, A. Seve, L. Courtellemont, H. Péré, L. Hocqueloux, S. Fafi-Kremer, T. Prazuck, H. Mouquet, T. Bruel, E. Simon-Lorière, F. A. Rey, O. Schwartz, Reduced sensitivity of SARS-CoV-2 variant Delta to antibody neutralization. Nature 596, 276–280 (2021).

13. D. J. Sheward, C. Kim, R. A. Ehling, A. Pankow, X. Castro Dopico, R. Dyrdak, D. P. Martin, S. T. Reddy, J. Dillner, G. B. Karlsson Hedestam, J. Albert, B. Murrell, Neutralisation sensitivity of the SARS-CoV-2 omicron (B.1.1.529) variant: a cross-sectional study. Lancet Infect. Dis. 22, 813–820 (2022).

14. L. Hanke, L. Vidakovics Perez, D. J. Sheward, H. Das, T. Schulte, A. Moliner-Morro, M. Corcoran, A. Achour, G. B. Karlsson Hedestam, B. M. Hällberg, B. Murrell, G. M. McInerney, An alpaca nanobody neutralizes SARS-CoV-2 by blocking receptor interaction. Nat. Commun. 11, 4420 (2020).

15. F. Muecksch, Y. Weisblum, C. O. Barnes, F. Schmidt, D. Schaefer-Babajew, Z. Wang, J. C. C Lorenzi, A. I. Flyak, A. T. DeLaitsch, K. E. Huey-Tubman, S. Hou, C. A. Schiffer, C. Gaebler, J. Da Silva, D. Poston, S. Finkin, A. Cho, M. Cipolla, T. Y. Oliveira, K. G. Millard, V. Ramos, A. Gazumyan, M. Rutkowska, M. Caskey, M. C. Nussenzweig, P. J. Bjorkman, T. Hatziioannou, P. D. Bieniasz, Affinity maturation of SARS-CoV-2 neutralizing antibodies confers potency, breadth, and resilience to viral escape mutations. Immunity 54, 1853–1868.e7 (2021).

16. D. J. Sheward, P. Pushparaj, H. Das, C. Kim, S. Kim, L. Hanke, R. Dyrdak, G. McInerney, J. Albert, B. Murrell, G. B. Karlsson Hedestam, B. Martin Hällberg, Structural basis of Omicron neutralization by affinity-matured public antibodies, bioRxiv (2022)p. 2022.01.03.474825.

17. J. M. Taft, C. R. Weber, B. Gao, R. A. Ehling, J. Han, L. Frei, S. W. Metcalfe, M. Overath, A. Yermanos, W. Kelton, S. T. Reddy, Deep Mutational Learning Predicts ACE2 Binding and Antibody Escape to Combinatorial Mutations in the SARS-CoV-2 Receptor Binding Domain. Cell, doi: 10.1016/j.cell.2022.08.024 (2022).

18. A. J. Greaney, T. N. Starr, C. O. Barnes, Y. Weisblum, F. Schmidt, M. Caskey, C. Gaebler, A. Cho, M. Agudelo, S. Finkin, Z. Wang, D. Poston, F. Muecksch, T. Hatziioannou, P. D. Bieniasz, D. F. Robbiani, M. C. Nussenzweig, P. J. Bjorkman, J. D. Bloom, Mapping mutations to the SARS-CoV-2 RBD that escape binding by different classes of antibodies. Nat. Commun. 12, 4196 (2021).

19. J. Zahradník, J. Nunvar, G. Schreiber, Perspectives: SARS-CoV-2 Spike Convergent Evolution as a Guide to Explore Adaptive Advantage. Front. Cell. Infect. Microbiol. 12, 748948 (2022).

20. D. P. Martin, S. Weaver, H. Tegally, J. E. San, S. D. Shank, E. Wilkinson, A. G. Lucaci, J. Giandhari, S. Naidoo, Y. Pillay, L. Singh, R. J. Lessells, NGS-SA, COVID-19 Genomics UK (COG-UK), R. K. Gupta, J. O. Wertheim, A. Nekturenko, B. Murrell, G. W. Harkins, P. Lemey, O. A. MacLean, D. L. Robertson, T. de Oliveira, S. L. Kosakovsky Pond, The emergence and ongoing convergent evolution of the SARS-CoV-2 N501Y lineages. Cell 184, 5189–5200.e7 (2021).

21. W. T. Harvey, A. M. Carabelli, B. Jackson, R. K. Gupta, E. C. Thomson, E. M. Harrison, C. Ludden, R. Reeve, A. Rambaut, COVID-19 Genomics UK (COG-UK) Consortium, S. J. Peacock, D. L. Robertson, SARS-CoV-2 variants, spike mutations and immune escape. Nat. Rev. Microbiol. 19, 409–424 (2021).

22. Y. Cao, F. Jian, J. Wang, Y. Yu, W. Song, A. Yisimayi, J. Wang, R. An, X. Chen, N. Zhang, Y. Wang, P. Wang, L. Zhao, H. Sun, L. Yu, S. Yang, X. Niu, T. Xiao, Q. Gu, F. Shao, X. Hao, Y. Xu, R. Jin, Z. Shen, Y. Wang, X. S. Xie, Imprinted SARS-CoV-2 humoral immunity induces convergent Omicron RBD evolution. Nature 614, 521–529 (2023).

23. W. Ma, H. Fu, F. Jian, Y. Cao, M. Li, Immune evasion and ACE2 binding affinity contribute to SARS-CoV-2 evolution. Nat Ecol Evol 7, 1457–1466 (2023).

24. T. N. Starr, A. J. Greaney, A. S. Dingens, J. D. Bloom, Complete map of SARS-CoV-2 RBD mutations that escape the monoclonal antibody LY-CoV555 and its cocktail with LY-CoV016. Cell Rep Med, 100255 (2021).

25. T. N. Starr, A. J. Greaney, A. Addetia, W. W. Hannon, M. C. Choudhary, A. S. Dingens, J. Z. Li, J. D. Bloom, Prospective mapping of viral mutations that escape antibodies used to treat COVID-19. Science 371, 850–854 (2021).

26. Z. Lin, H. Akin, R. Rao, B. Hie, Z. Zhu, W. Lu, N. Smetanin, R. Verkuil, O. Kabeli, Y. Shmueli, A. Dos Santos Costa, M. Fazel-Zarandi, T. Sercu, S. Candido, A. Rives, Evolutionary-scale prediction of atomic-level protein structure with a language model. Science 379, 1123–1130 (2023).

27. J. Lanchantin, T. Wang, V. Ordonez, Y. Qi, “General multi-label image classification with transformers” in 2021 IEEE/CVF Conference on Computer Vision and Pattern Recognition (CVPR) (IEEE, 2021; https://openaccess.thecvf.com/content/CVPR2021/html/Lanchantin_General_Multi-Label_Image_Classification_With_Transformers_CVPR_2021_paper.html).

28. K. Uriu, J. Ito, Y. Kosugi, Y. L. Tanaka, Y. Mugita, Z. Guo, A. A. Hinay, O. Putri, Y. Kim, R. Shimizu, M. M. Begum, M. Jonathan, A. Saito, T. Ikeda, K. Sato, Transmissibility, infectivity, and immune evasion of the SARS-CoV-2 BA.2.86 variant. Lancet Infect. Dis. 0 (2023).

29. S. Yang, Y. Yu, Y. Xu, F. Jian, W. Song, A. Yisimayi, P. Wang, J. Wang, J. Liu, L. Yu, X. Niu, J. Wang, Y. Wang, F. Shao, R. Jin, Y. Wang, Y. Cao, Fast evolution of SARS-CoV-2 BA.2.86 to JN.1 under heavy immune pressure. Lancet Infect. Dis. 24, e70–e72 (2024).

30. D. Chicco, G. Jurman, The advantages of the Matthews correlation coefficient (MCC) over F1 score and accuracy in binary classification evaluation. BMC Genomics 21, 6 (2020).

31. Q. He, L. Wu, Z. Xu, X. Wang, Y. Xie, Y. Chai, A. Zheng, J. Zhou, S. Qiao, M. Huang, G. Shang, X. Zhao, Y. Feng, J. Qi, G. F. Gao, Q. Wang, An updated atlas of antibody evasion by SARS-CoV-2 Omicron sub-variants including BQ.1.1 and XBB. Cell Rep Med 4, 100991 (2023).

32. J. Vig, A. Madani, L. R. Varshney, C. Xiong, R. Socher, N. Rajani, “BERTology Meets Biology: Interpreting Attention in Protein Language Models” in International Conference on Learning Representations (2021; https://openreview.net/forum?id=YWtLZvLmud7).

33. S. Kumar, K. Karuppanan, G. Subramaniam, Omicron (BA.1) and sub-variants (BA.1.1, BA.2, and BA.3) of SARS-CoV-2 spike infectivity and pathogenicity: A comparative sequence and structural-based computational assessment. J. Med. Virol. 94, 4780–4791 (2022).

34. Z. Liu, L. A. VanBlargan, L.-M. Bloyet, P. W. Rothlauf, R. E. Chen, S. Stumpf, H. Zhao, J. M. Errico, E. S. Theel, M. J. Liebeskind, B. Alford, W. J. Buchser, A. H. Ellebedy, D. H. Fremont, M. S. Diamond, S. P. J. Whelan, Identification of SARS-CoV-2 spike mutations that attenuate monoclonal and serum antibody neutralization. Cell Host Microbe 29, 477–488.e4 (2021).

35. L. Frei, B. Gao, J. Han, J. M. Taft, E. B. Irvine, C. R. Weber, R. K. Kumar, B. N. Eisinger, S. T. Reddy, Deep learning-guided selection of antibody therapies with enhanced resistance to current and prospective SARS-CoV-2 Omicron variants, bioRxiv (2023)p. 2023.10.09.561492.

36. T. Kuramochi, S. W. Gan, A. W. S. Ho, B. Wang, N. Kageji, T. Nambu, S. Iida, M. Okuda-Miura, W. S. Chia, C. Y. Yeo, D. Chen, W.-H. Lee, E. Z. X. Ngoh, S. N. Mohd Salleh, C.-I. Wang, T. Igawa, H. Shimada, Comprehensive engineering of a therapeutic neutralizing antibody targeting SARS-CoV-2 spike protein to neutralize escape variants. MAbs 14, 2040350 (2022).

37. C. G. Rappazzo, L. V. Tse, C. I. Kaku, D. Wrapp, M. Sakharkar, D. Huang, L. M. Deveau, T. J. Yockachonis, A. S. Herbert, M. B. Battles, C. M. O’Brien, M. E. Brown, J. C. Geoghegan, J. Belk, L. Peng, L. Yang, Y. Hou, T. D. Scobey, D. R. Burton, D. Nemazee, J. M. Dye, J. E. Voss, B. M. Gunn, J. S. McLellan, R. S. Baric, L. E. Gralinski, L. M. Walker, Broad and potent activity against SARS-like viruses by an engineered human monoclonal antibody. Science 371, 823–829 (2021).

38. M. G. Ison, D. F. Weinstein, M. Dobryanska, A. Holmes, A.-M. Phelan, Y. Li, D. Gupta, K. Narayan, K. Tosh, E. Hershberger, L. E. Connolly, I. Yalcin, E. Campanaro, P. Hawn, P. Schmidt, EVADE Study Group, Prevention of COVID-19 Following a Single Intramuscular Administration of Adintrevimab: Results From a Phase 2/3 Randomized, Double-Blind, Placebo-Controlled Trial (EVADE). Open Forum Infect Dis 10, ofad314 (2023).

39. B. West, A. Z. Wec, M. Doyle, C. Kaku, P. Hawn, L. Dillinger, L. Walker, “NVD200 potently neutralises Omicron and its sublineages” in 33rd European Congress of Clinical Microbiology and Infectious Diseases (2023; https://invivyd.com/wp-content/uploads/2023/08/ECCMID-2023_NVD200-potently-neutralises-Omicron-and-its-sublineages-_Final-Poster.pdf).

40. Center for Drug Evaluation, Research, Emergency Use Authorizations for Drugs and Non-Vaccine Biological Products, U.S. Food and Drug Administration (2024). https://www.fda.gov/drugs/emergency-preparedness-drugs/emergency-use-authorizations-drugs-and-non-vaccine-biological-products.

41. N. Houlsby, A. Giurgiu, S. Jastrzebski, B. Morrone, Q. De Laroussilhe, A. Gesmundo, M. Attariyan, S. Gelly, “Parameter-Efficient Transfer Learning for NLP” in Proceedings of the 36th International Conference on Machine Learning, K. Chaudhuri, R. Salakhutdinov, Eds. (PMLR, 09--15 Jun 2019; https://proceedings.mlr.press/v97/houlsby19a.html)vol. 97 of Proceedings of Machine Learning Research, pp. 2790–2799.

42. H. Liu, D. Tam, M. Muqeeth, J. Mohta, T. Huang, M. Bansal, C. Raffel, Few-shot parameter-efficient fine-tuning is better and cheaper than in-context learning, S. Koyejo, S. Mohamed, A. Agarwal, D. Belgrave, K. Cho, A. Oh, Eds., arXiv [cs.LG] (2022)pp. 1950–1965.

43. Y. Mao, L. Mathias, R. Hou, A. Almahairi, H. Ma, J. Han, S. Yih, M. Khabsai, “UniPELT: A Unified Framework for Parameter-Efficient Language Model Tuning” in Proceedings of the 60th Annual Meeting of the Association for Computational Linguistics (Volume 1: Long Papers), S. Muresan, P. Nakov, A. Villavicencio, Eds. (Association for Computational Linguistics, Dublin, Ireland, 2022; https://aclanthology.org/2022.acl-long.433), pp. 6253–6264.

44. R. Vazquez-Lombardi, D. Nevoltris, A. Luthra, P. Schofield, C. Zimmermann, D. Christ, Transient expression of human antibodies in mammalian cells. Nat. Protoc. 13, 99–117 (2018).

45. E. T. Boder, K. D. Wittrup, Yeast surface display for screening combinatorial polypeptide libraries. Nat. Biotechnol. 15, 553–557 (1997).

46. D. J. Sheward, M. Mandolesi, E. Urgard, C. Kim, L. Hanke, L. P. Vidakovics, A. Pankow, N. L. Smith, X. C. Dopico, G. M. McInerney, Others, Beta RBD boost broadens antibody-mediated protection against SARS-CoV-2 variants in animal models. Cell Reports Medicine 2, 100450 (2021).

47. D. M. Mason, S. Friedensohn, C. R. Weber, C. Jordi, B. Wagner, S. M. Meng, R. A. Ehling, L. Bonati, J. Dahinden, P. Gainza, B. E. Correia, S. T. Reddy, Optimization of therapeutic antibodies by predicting antigen specificity from antibody sequence via deep learning. Nat Biomed Eng 5, 600–612 (2021).

48. R Core Team, R: A Language and Environment for Statistical Computing. R Foundation for Statistical Computing [Preprint] (2021). https://www.R-project.org/.

49. G. Van Rossum, F. L. Drake, Python 3 Reference Manual: (Python Documentation Manual Part 2) (CreateSpace Independent Publishing Platform, 2009; https://play.google.com/store/books/details?id=KIybQQAACAAJ).

50. H. Wickham, ggplot2: Elegant Graphics for Data Analysis. Springer-Verlag New York [Preprint] (2016). https://ggplot2.tidyverse.org.

51. J. D. Hunter, Matplotlib: A 2D Graphics Environment. Comput. Sci. Eng. 9, 90–95 (May-June 2007).

52. Z. Gu, R. Eils, M. Schlesner, Complex heatmaps reveal patterns and correlations in multidimensional genomic data. [Preprint] (2016).

53. R. Kolde, Others, Pheatmap: pretty heatmaps. R package version.

54. G. Csardi, T. Nepusz, The igraph software package for complex network research. researchgate.net.

55. J. A. Gustavsen, S. Pai, R. Isserlin, B. Demchak, A. R. Pico, RCy3: Network biology using Cytoscape from within R. F1000Res. 8, 1774 (2019).

56. H. Wickham, stringr: Simple, Consistent Wrappers for Common String Operations. [Preprint] (2023). https://stringr.tidyverse.org.

57. H. Wickham, R. François, L. Henry, K. Müller, dplyr: A Grammar of Data Manipulation. [Preprint] (2022).

58. E. Neuwirth, RColorBrewer: ColorBrewer palettes. R package version 1.1-2. (2014).

59. The pandas development team, Pandas-Dev/pandas: Pandas (Zenodo, 2024; 10.5281/zenodo.3509134).

60. C. R. Harris, K. J. Millman, S. J. van der Walt, R. Gommers, P. Virtanen, D. Cournapeau, E. Wieser, J. Taylor, S. Berg, N. J. Smith, R. Kern, M. Picus, S. Hoyer, M. H. van Kerkwijk, M. Brett, A. Haldane, J. F. Del Río, M. Wiebe, P. Peterson, P. Gérard-Marchant, K. Sheppard, T. Reddy, W. Weckesser, H. Abbasi, C. Gohlke, T. E. Oliphant, Array programming with NumPy. Nature 585, 357–362 (2020).

61. K. Sechidis, G. Tsoumakas, I. Vlahavas, “On the Stratification of Multi-label Data” in Machine Learning and Knowledge Discovery in Databases (Springer Berlin Heidelberg, 2011; 10.1007/978-3-642-23808-6_10), pp. 145–158.

62. J. L. Ba, J. R. Kiros, G. E. Hinton, Layer Normalization, *arXiv [stat.ML]* (2016). http://arxiv.org/abs/1607.06450.

63. Chen, Xu, Hui, Wu, Lin, “Learning Semantic-Specific Graph Representation for Multi-Label Image Recognition” in 2019 IEEE/CVF International Conference on Computer Vision (ICCV) (2019; 10.1109/ICCV.2019.00061)vol. 0, pp. 522–531.

64. A. Paszke, S. Gross, F. Massa, A. Lerer, J. Bradbury, G. Chanan, T. Killeen, Z. Lin, N. Gimelshein, L. Antiga, A. Desmaison, A. Köpf, E. Yang, Z. DeVito, M. Raison, A. Tejani, S. Chilamkurthy, B. Steiner, L. Fang, J. Bai, S. Chintala, “PyTorch: an imperative style, high-performance deep learning library” in *Proceedings of the 33rd International Conference on Neural Information Processing Systems* (Curran Associates Inc., Red Hook, NY, USA, 2019), pp. 8026–8037.

65. S. Cook, CUDA Programming: A Developer’s Guide to Parallel Computing with GPUs (Newnes, 2012; https://play.google.com/store/books/details?id=g3EzsZn4poUC).

66. Y. LeCun, Y. Bengio, “Convolutional networks for images, speech, and time series” in The Handbook of Brain Theory and Neural Networks (MIT Press, Cambridge, MA, USA, 1998), pp. 255–258.

