## Supplementary material for "Synthetic coevolution reveals adaptive mutational trajectories of neutralizing antibodies and SARS-CoV-2": Suppplementary_Materials_RE_MM_SynCoevo.pdf

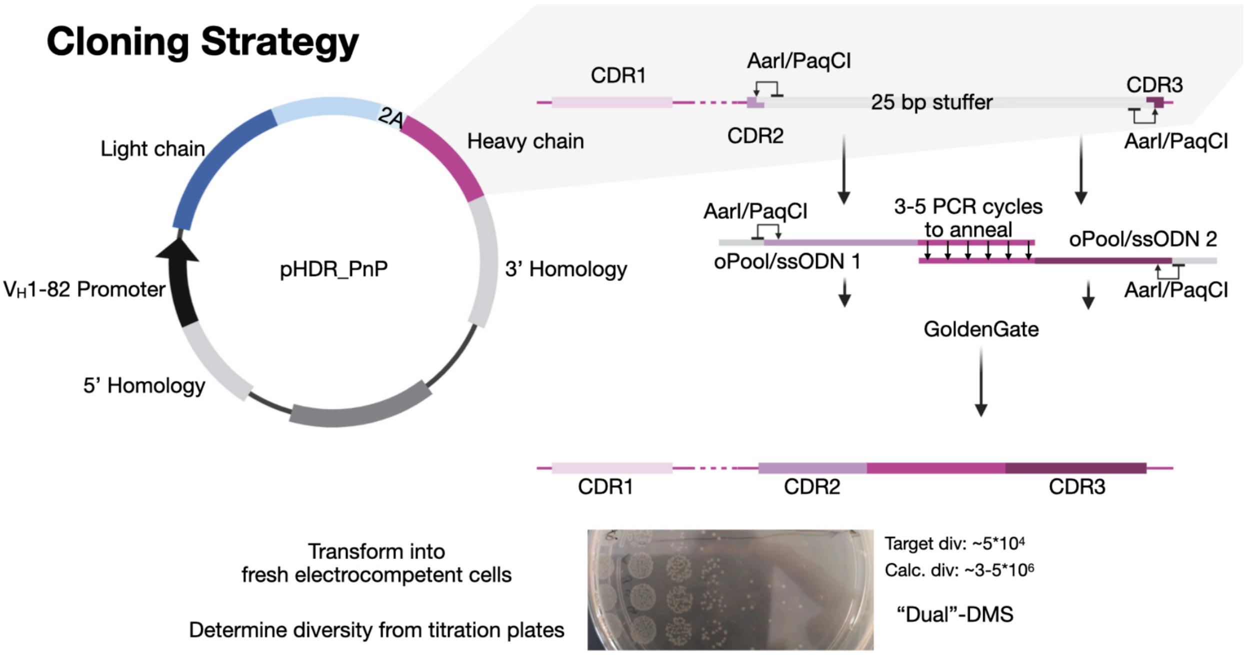


**Figure S1. Genetic assembly and cloning of antibody clonotype libraries incorporating synthetic somatic hypermutation.** Golden gate cloning and PCR is used to assemble an entry vector comprising a stuffer region flanked by restriction sites for PaqCI and single-stranded oligonucleotides (ssODN). The ssODNs consist of a pool (oPool) of tiled NNK/NNN mutations targeted to antibody variable heavy chain (VH) complementarity determining region 2 (HCDR2) and HDCR3. The Golden Gate assembly product is transformed into fresh electrocompetent cells. Titration plates are used to approximate the library size. Midiprepped plasmid is then used as a homology-directed repair (HDR) template CRISPR-Cas9-mediated integration into a mammalian antibody display and secretion platform (plug-and-(dis)play, PnP cells).

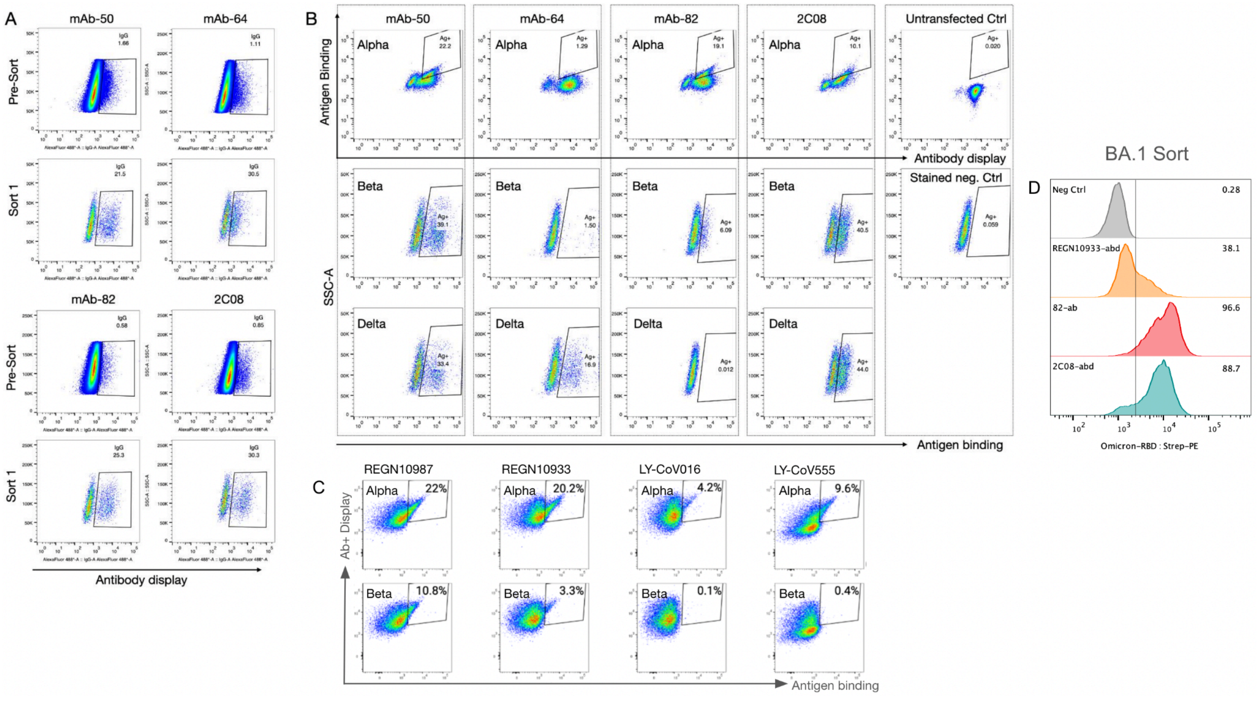


**Figure S2. Mammalian antibody display and screening of clonotype libraries for binding to RBD variants.** (**A**) Following CRISPR-Cas9-mediated HDR integration of sSHM (antibody clonotype) libraries into PnP cells, FACS was performed to isolate PnP cells expressing functional surface antibody. (**B, C**) The PnP antibody clonotype (sSHM) libraries were sorted for binding to SARS-CoV-2 RBD variants Alpha, Beta or Delta (B); or Alpha or Beta (C). (**D**) Histograms show Alpha, Beta and Delta RBD (pooled)-selected antibody clonotype libraries and their binding to BA.1 RBD (results are following two rounds FACS enrichment).

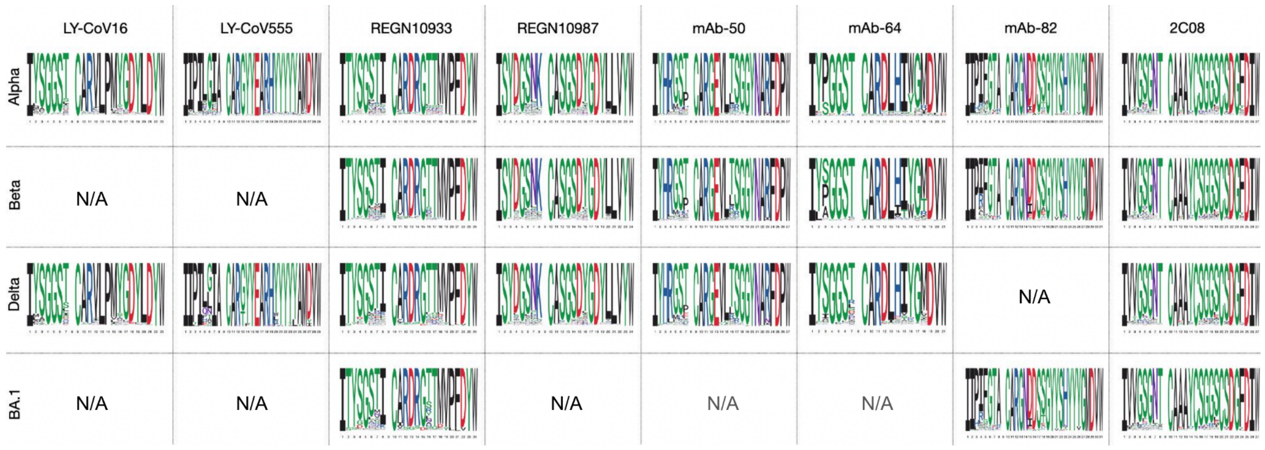


**Figure S3. Sequence diversity of antibody clonotype (sSHM) libraries following RBD-variant selection.** Amino acid sequence logos derived from targeted deep sequencing of the antibody clonotype libraries post-selection of binding to RBD variants. Depicted are the HCDR2 and HCDR3 of the various antibodies.

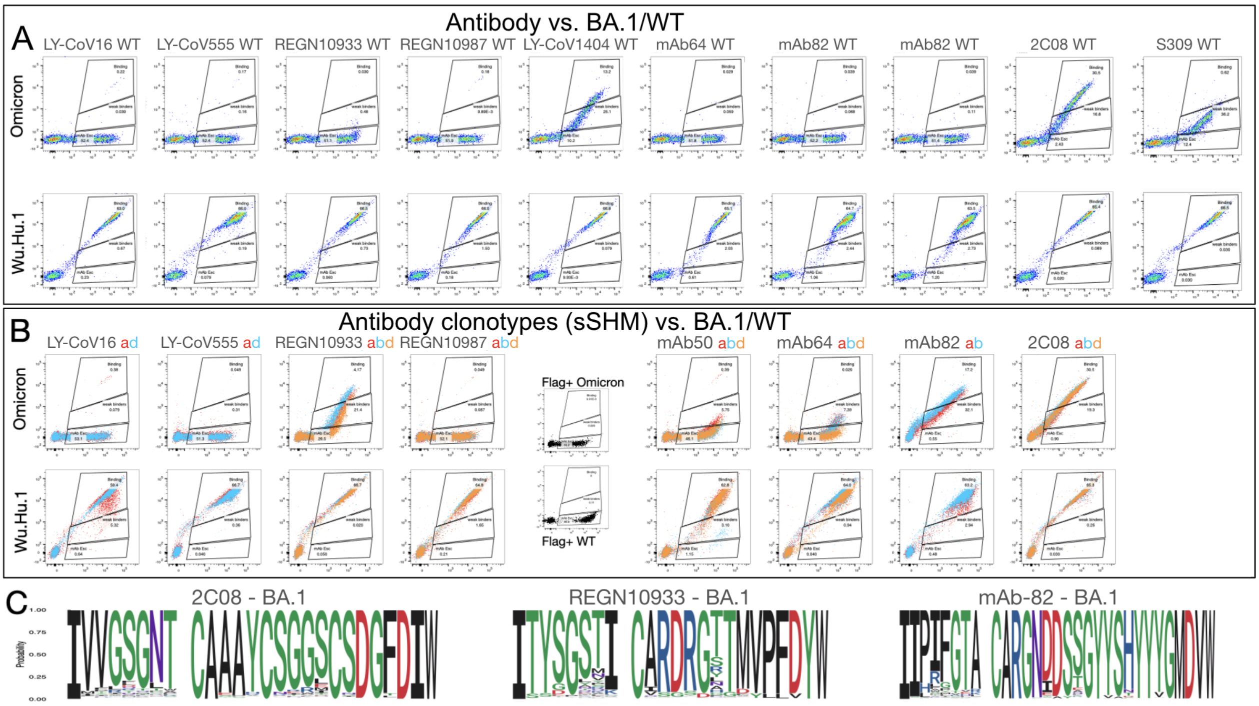


**Figure S4. Omicron reactivity of antibodies and antibody libraries.** (**A, B**) Omicron BA.1 (top row) and ancestral Wu-Hu-1 RBD (bottom row) expressed by yeast surface display are screened for binding or escape to wild-type antibodies (A) or Alpha (a), Beta (b), Delta (d) variant-selected antibody clonotype libraries (B). Flag-only control is shown in middle (black dot plot) (B). (**C**). Amino acid sequence logos derived from targeted deep sequencing of the antibody clonotype libraries post-selection of binding to BA.1 RBD. Depicted are the HCDR2 and HCDR3 of the various antibodies.

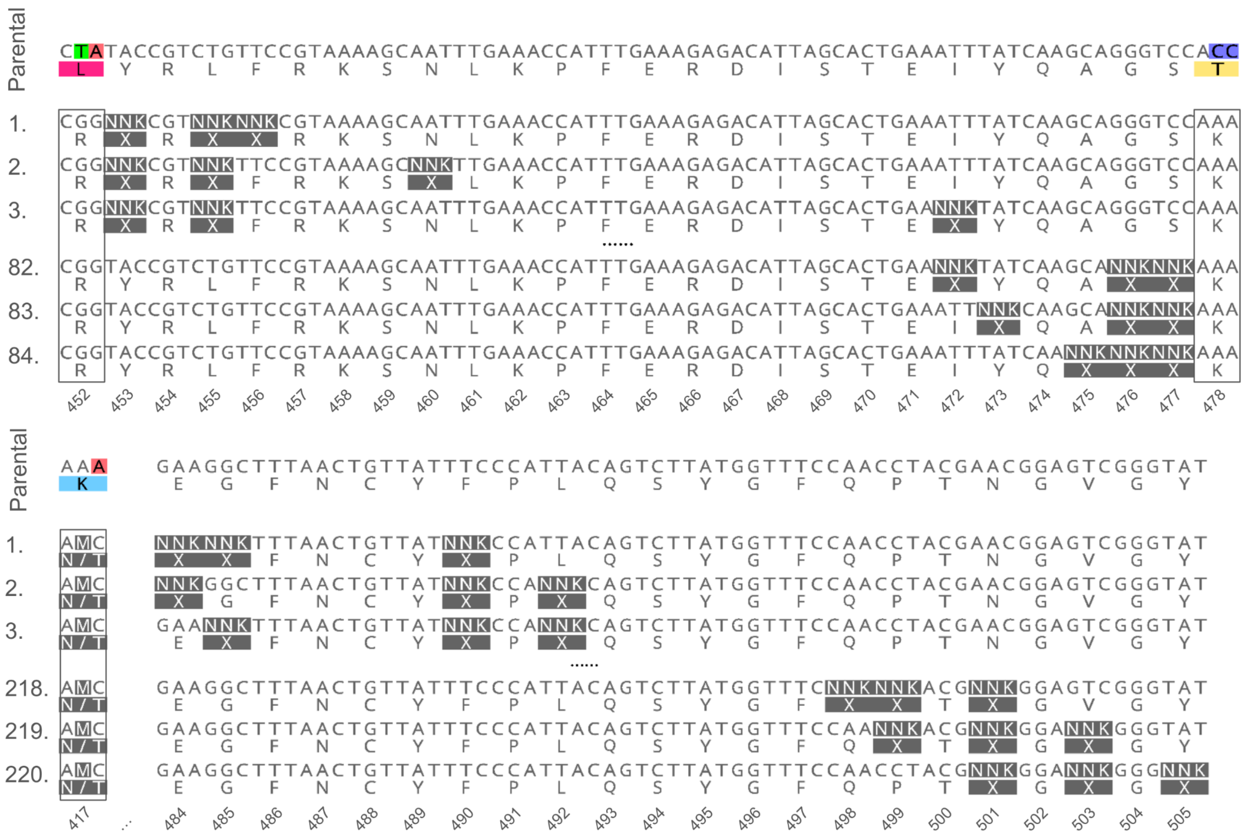


**Figure S5. The design of extended RBD variant libraries with additional tiled NNK mutagenesis.** Top: library RBM-1$_{L452R+T478K}$, which consists of three tiled NNK codons across RBM-1 and containing the frequently observed mutations L452R (Delta, BA.4/5) and T478K (Delta, BA.1/2/3/4/5). Bottom: library RBM-2$_{K417N/T}$, which consists of three tiled NNK codons across RBM-2 and possessing the dominant K417N and observed K417T mutation.

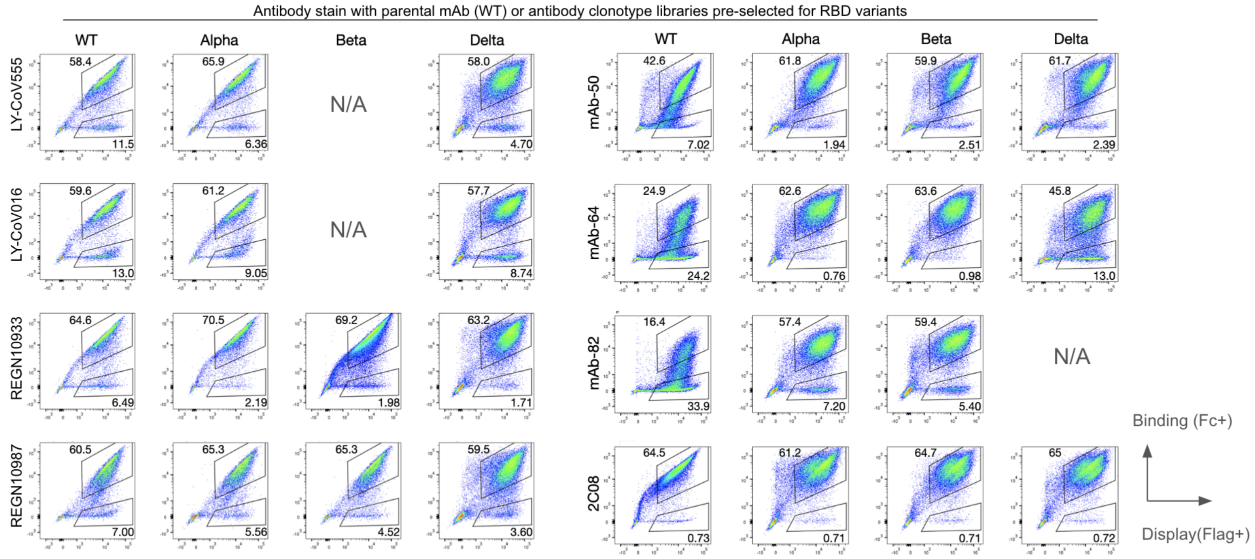


**Figure S6. Antibody and clonotype pools screened for binding or escape to RBD variants expressed by yeast surface display.** The parental antibody (WT) or its clonotype (sSHM) libraries were pre-selected for binding to RBD variants (Alpha, Beta or Delta); shown are flow cytometry dot plots of their binding or escape to yeast display of RBD variants (pooled libraries: RBM-1T, RBM-2T, RBM-3T).

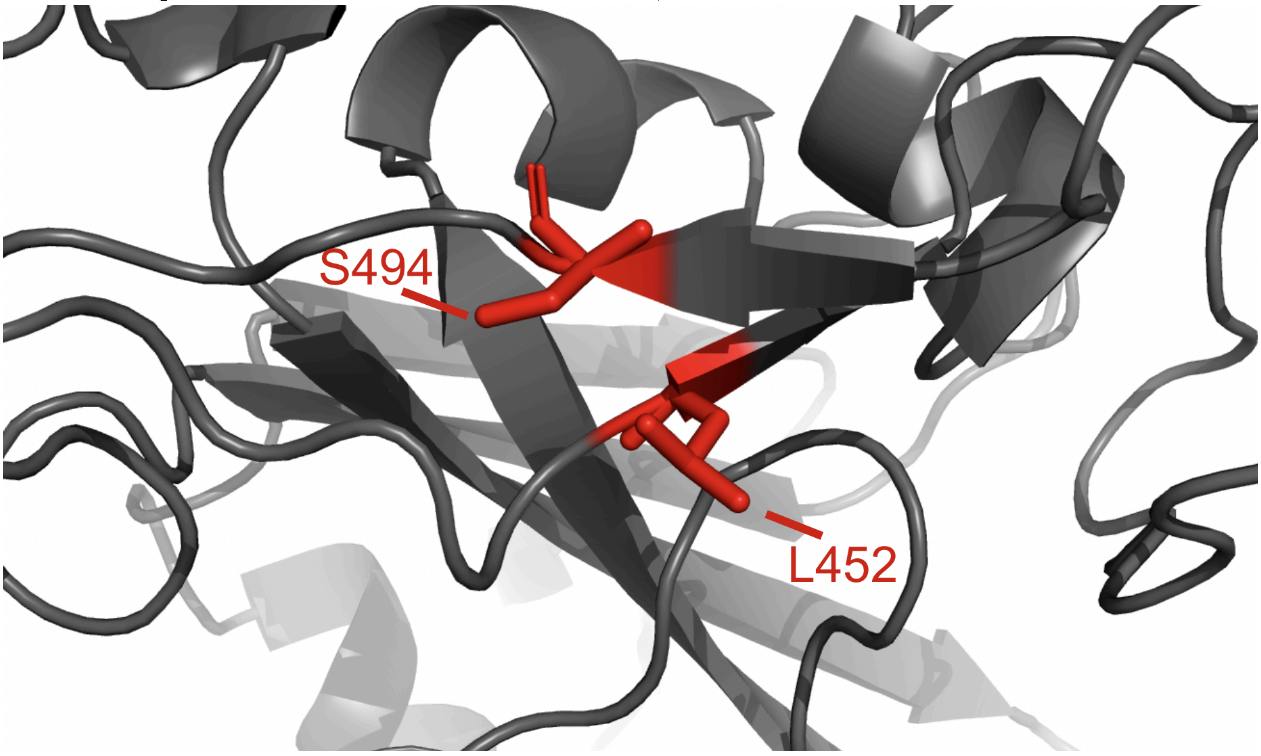


**Figure S7. Structural analysis of co-occurring mutations in the RBD.** Proximity of S494 and L452 (red) within the RBD (dark gray**)**.

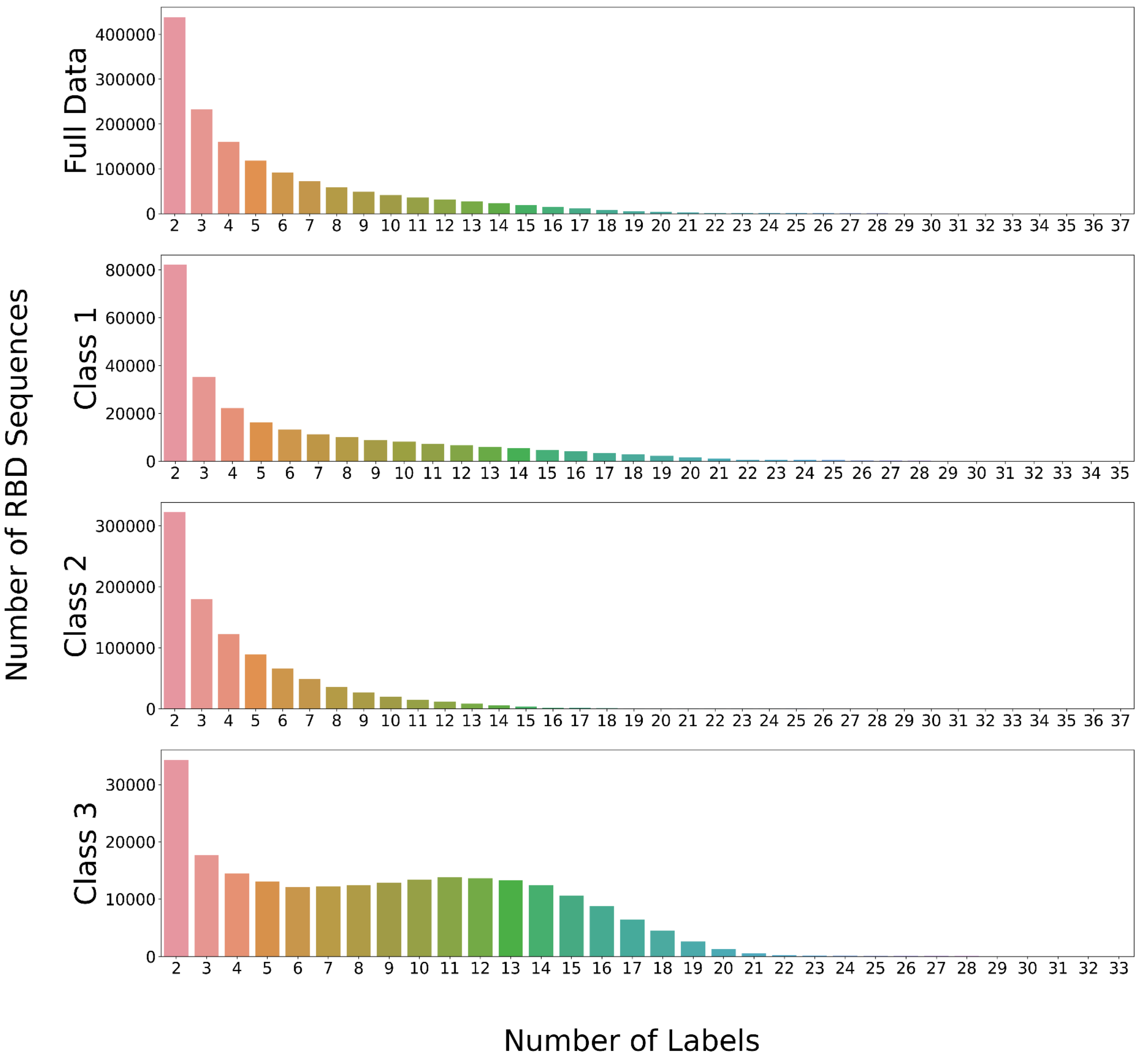


**Figure S8. Distribution of antibody label frequency per RBM class**. Histograms representing the distribution of the number of antibody (and ACE2) labels for RBD variants. Label distribution is presented for the full dataset as well as the individual RBM-1, -2, and -3.

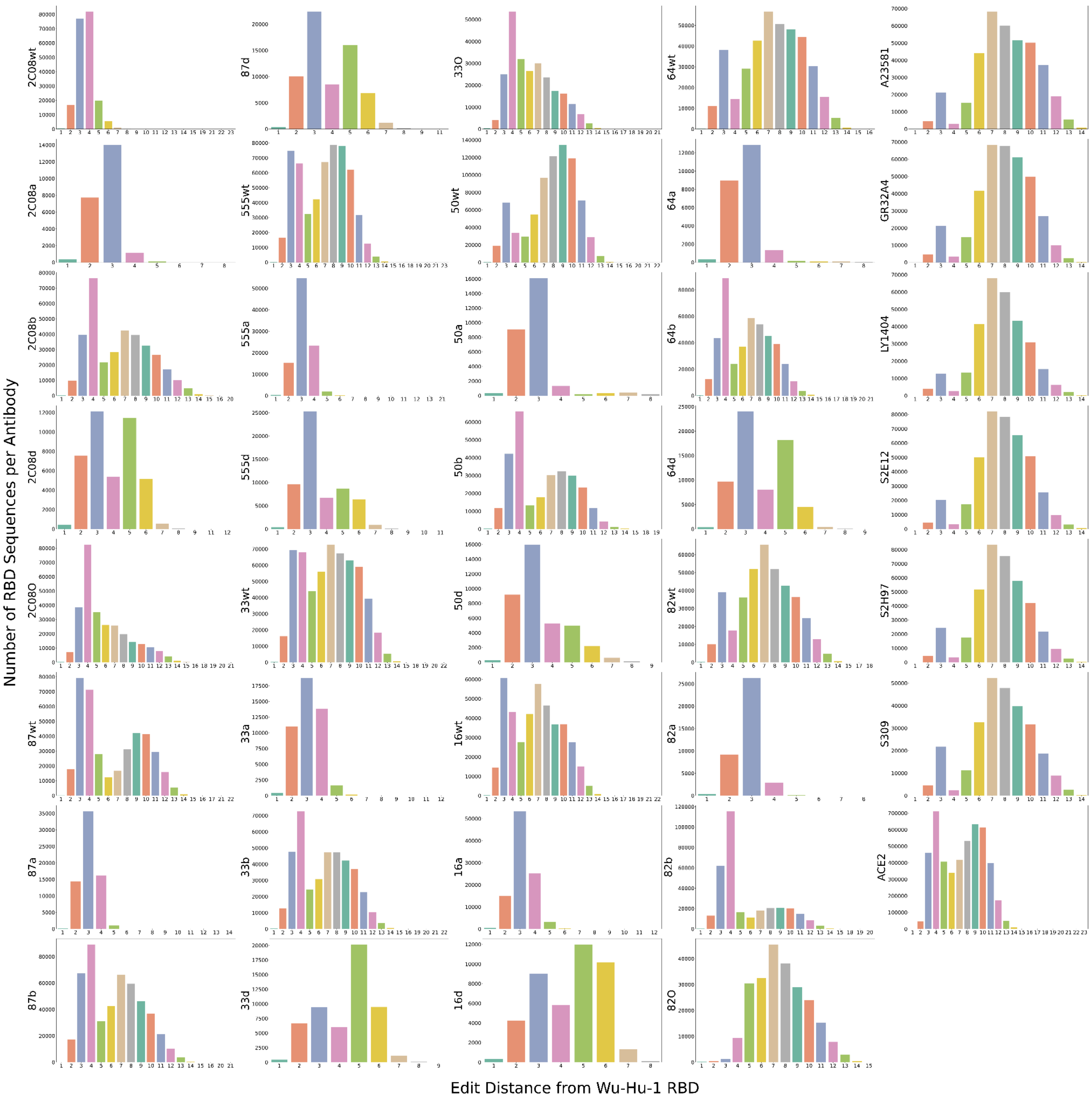


**Figure S9. Edit distance of RBD variant sequences from ancestral Wu-Hu-1 RBD sequence**. Individual histograms represent the frequency of different edit distances (a.a.) of RBD variants recovered following antibody or ACE2 binding and escape selections. Edit distance is calculated from the reference sequence of the Wu-Hu-1 RBD. Antibody abbreviations: 16: LY-CoV16, 33: REGN10933, 87: REGN10987, 555: LY-CoV555. Variant abbreviations: a: Alpha, b: Beta, d: Delta, O: Omicron. sSHM antibody nomenclature indicates the selection variant e.g. 16d refers to LY-CoV16 selected on Delta. WT refers to wild-type antibodies.

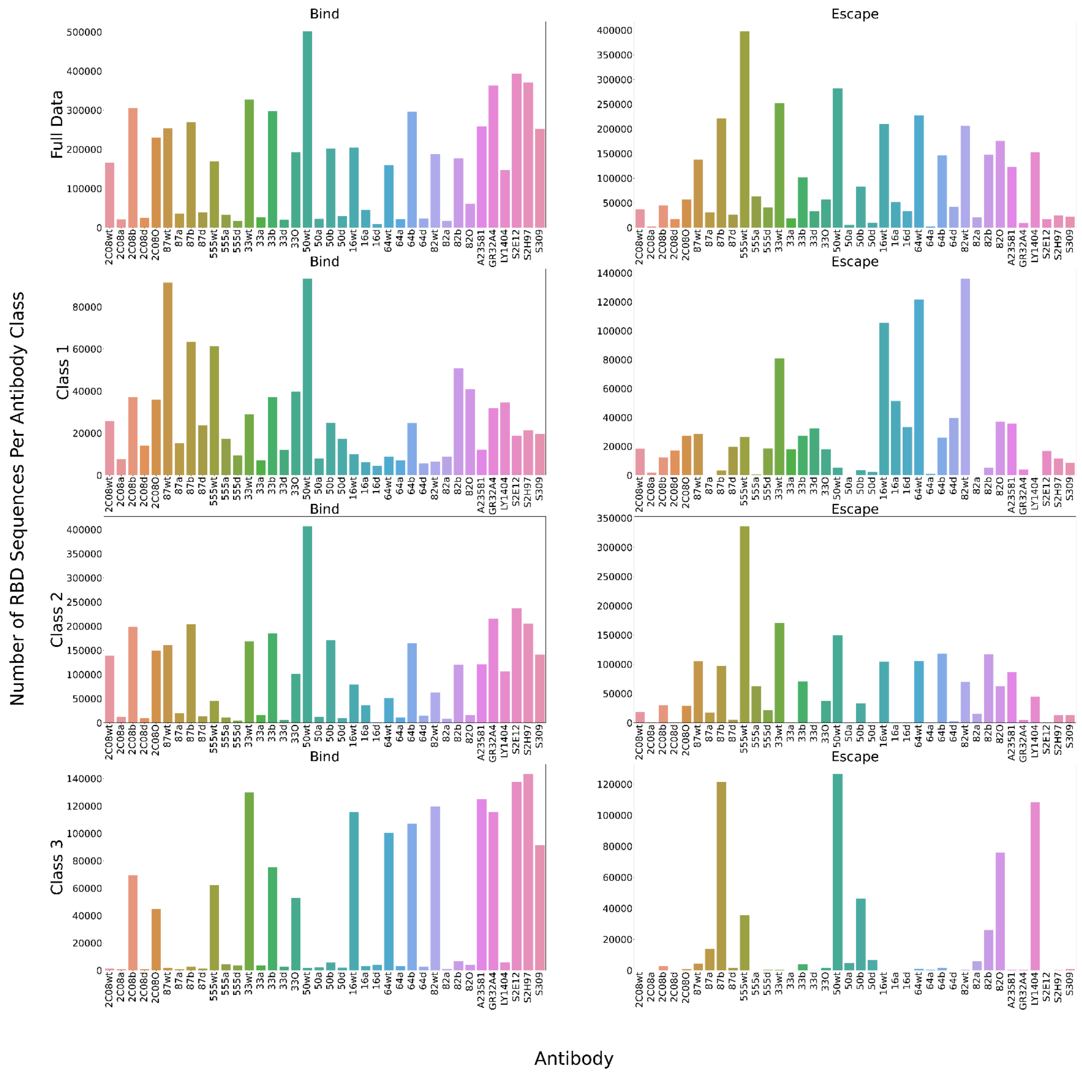


**Figure S10. Number of RBD variant sequences per RBM class recovered after antibody selections**. The number of RBD variants recovered after selection for antibody binding or escape and stratified based on RBM-1, -2 and -3. Antibody abbreviations: 16: LY-CoV16, 33: REGN10933, 87: REGN10987, 555: LY-CoV555. Variant abbreviations: a: Alpha, b: Beta, d: Delta, O: Omicron. sSHM antibody nomenclature indicates the selection variant e.g. 16d refers to LY-CoV16 selected on Delta. WT refers to wild-type antibodies.

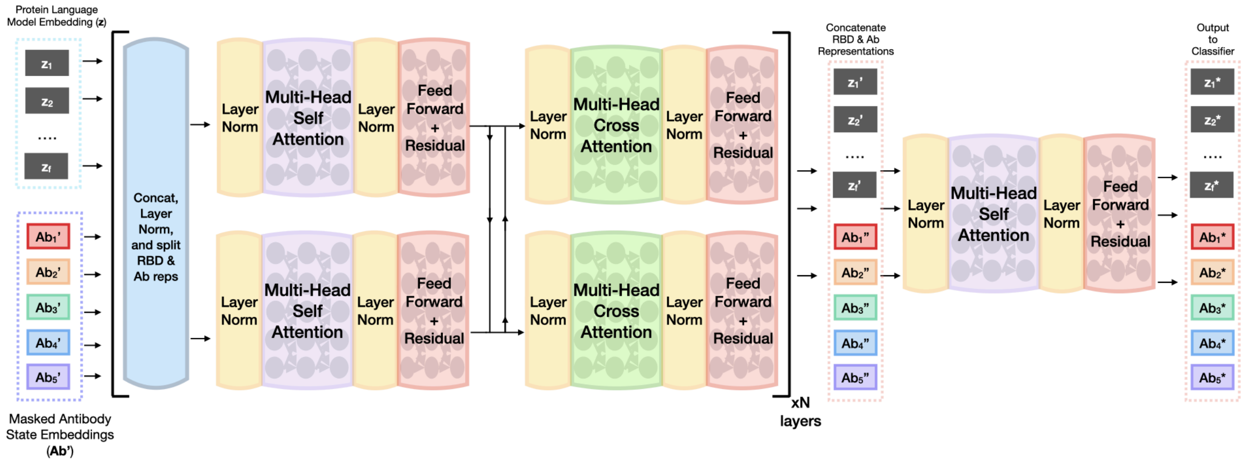


**Figure S11. RBD-pLM inter-attention mechanism**. RBD and antibody representations are fed to two inter-attention layers followed by concatenation of the RBD and antibody embeddings and a multi-headed self-attention block. Inter-attention layers consist of a layer normalization projection on the concatenated RBD and antibody embeddings. The output is then split back into RBD and antibody embeddings, which are then subjected to self-attention blocks in parallel, followed by cross-attention blocks.

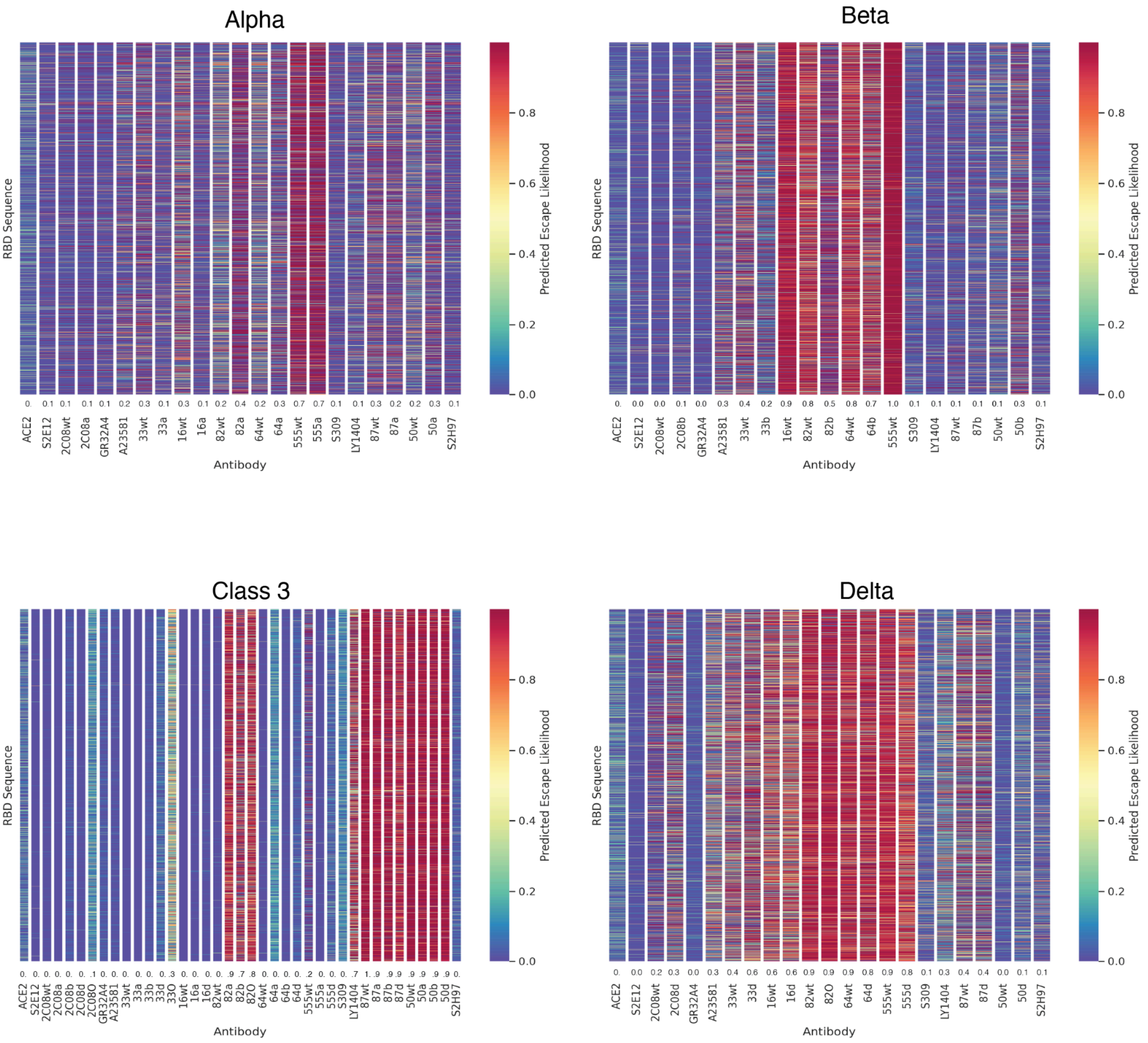


**Figure S12. Viral escape predictions of synthetic variant lineages**. Synthetic variant lineage RBD sequences for Alpha, Beta, Delta, and RBM-3 (Class 3) variants were generated as previously described (*17*, *35*). RBD-pLM escape predictions are shown for wild type antibodies and sSHM antibody pools selected for the parent variant of the synthetic lineage sequences (e.g. 16wt, 16d for Delta). The binding/escape prediction threshold is set to 0.5. Above 0.5 is considered escape (red). 0.5 or below is considered binding (yellow to blue). A summary statistic describing the total number of escaping RBD variants is divided by the total number of variants, calculated per target (antibodies and ACE2) and reported at the bottom of the columns. Antibody abbreviations: 16: LY-CoV16, 33: REGN10933, 87: REGN10987, 555: LY-CoV555. Variant abbreviations: a: Alpha, b: Beta, d: Delta, O: Omicron. sSHM antibody nomenclature indicates the selection variant e.g. 16d refers to LY-CoV16 selected on Delta. WT refers to wild-type antibodies.

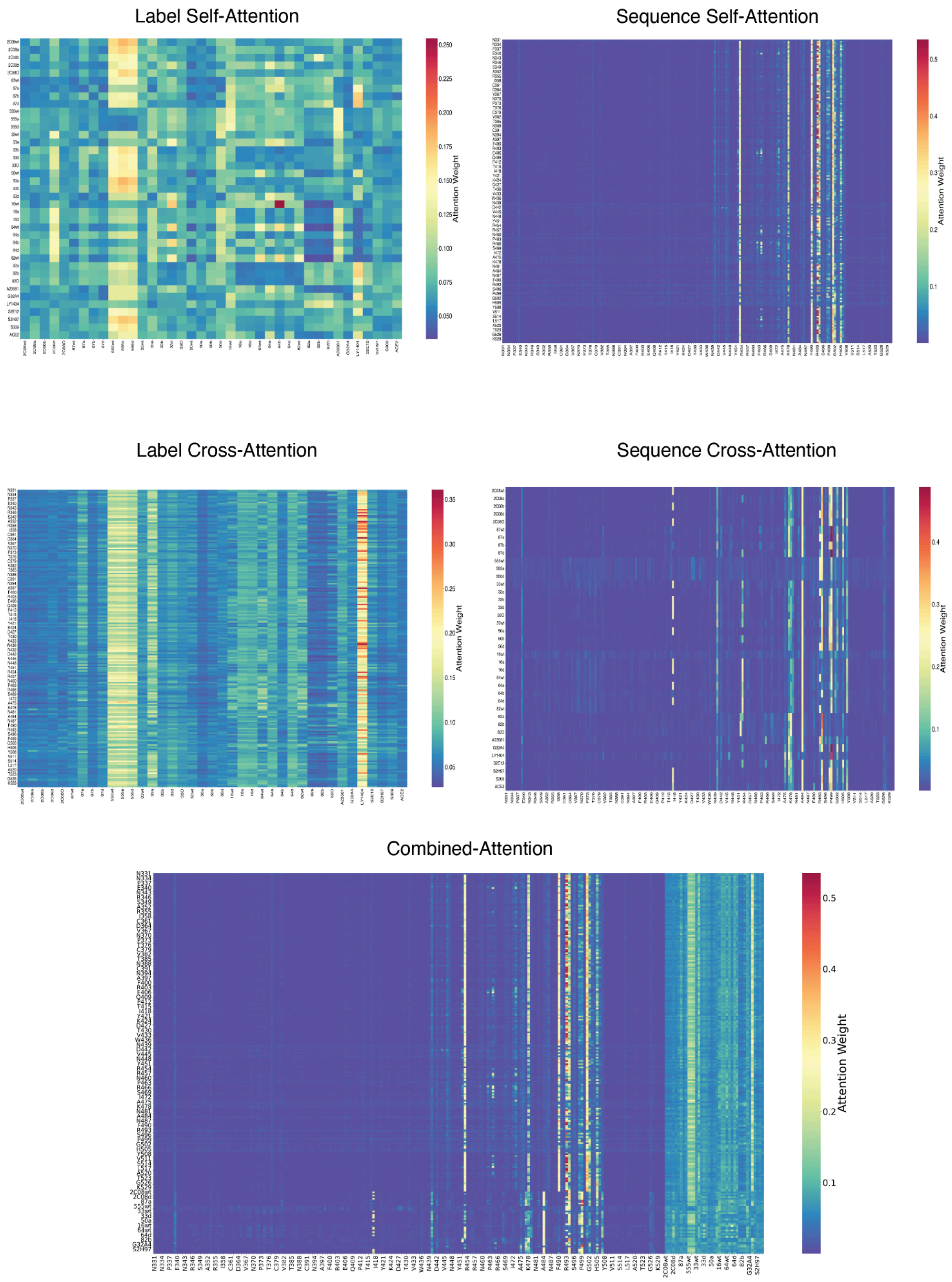


**Figure S13. RBD-pLM attention weights for variant BA1.1**. Attention weights are plotted for the five components of the inter-attention mechanism: (i) label self-attention, (ii) sequence self-attention, (iii) label cross-attention, (iv) sequence cross-attention and (v) combined-attention. For a given inter-attention layer and component, attention weights are averaged over all attention heads. Attentions for each component are then summed for all layers and the resulting matrix of weights is used to generate heatmaps. Note: combined-attention is a summation of all inter-attention components with the final inter-attention self-attention layer, in which sequence and label embeddings are concatenated together. Antibody abbreviations: 16: LY-CoV16, 33: REGN10933, 87: REGN10987, 555: LY-CoV555. Variant abbreviations: a: Alpha, b: Beta, d: Delta, O: Omicron. sSHM antibody nomenclature indicates the selection variant e.g. 16d refers to LY-CoV16 selected on Delta. WT refers to wild-type antibodies.

**Table S1. List of neutralizing antibodies used in this study for synthetic coevolution.**

| Antibody | Class | Description | Reference |
|---|---|---|---|
| LY-CoV16 (Etesevimab ) | 1 | Neutralizing antibody used as part of the Eli-Lilly therapeutic antibody cocktail. Derived from convalescent COVID-19 patient samples. LY-CoV16 is no longer authorized for clinical use due to loss of neutralization to Omicron BA.1 or sublineages. | Shi et al. Nature. 2020 |
| LY-CoV555 (Bamlanivimab) | 2 | Neutralizing antibody used as part of the Eli-Lilly therapeutic antibody cocktail. Identified through high-throughput screening of B cells from convalescent COVID-19 patient samples. LY-CoV555 is no longer authorized for clinical use due to loss of neutralization to Omicron BA.1 or sublineages. | Jones et al. Science Transl. Med. 2021 |
| REGN10933 (Casirivimab) | 1 | Neutralizing antibody used as part of the Regeneron COVID-19 therapeutic antibody cocktail. Derived from genetically humanized mice or convalescent human COVID-19 patient samples. REGN10933 is no longer authorized for clinical use due to loss of neutralization to Omicron BA.1 or sublineages. | Hansen et al. Science, 2020 |
| REGN10987 (Imdevimab) | 3 | Neutralizing antibody used as part of the Regeneron COVID-19 therapeutic antibody cocktail. Derived from genetically humanized mice or convalescent human COVID-19 patient samples. REGN10987 is no longer authorized for clinical use due to loss of neutralization to Omicron BA.1 or sublineages. | Hansen et al. Science, 2020 |
| mAb-50 | 3 | Neutralizing antibody isolated from expanded plasma cells of convalescent COVID-19 patients. | Ehling et al. Cell Rep, 2021. |
| mAb-64 | 1 | Neutralizing antibody isolated from expanded plasma cells of convalescent COVID-19 patients. | Ehling et al. Cell Rep, 2021. |
| mAb-82 | 1 | Neutralizing antibody isolated from expanded plasma cells of convalescent COVID-19 patients. | Ehling et al. Cell Rep, 2021. |
| A23-58.1 | 1 | Neutralizing antibody isolated from B cells of convalescent COVID-19 patients and shown to have broad neutralization. | Wang et al. Science, 2021 |
| 2C08 | 1 | Neutralizing antibody isolated from B cells of vaccinated COVID-19 patients and shown to have broad neutralization. Public clone induced by vaccination. | Schmitz et al. Cell, 2021 |

**Table S2. Dataset sizes per task**

| Dataset | Training Sequences | Validation Sequences | Test Sequences |
|---|---|---|---|
| ED 3 | 128,558 | 55,204 | 753,974 |
| ED 10 | 591,675 | 253,662 | 92,399 |
| Full Data | 651,274 | 133,238 | 158,224 |

**Table S3. Aggregate Machine Learning Metrics**
Due to its size, this table is included as a separate csv file

**Table S4. Model Performance (MCC) Per Antibody**
Due to its size, this table is included as a separate csv file
